# RNA-protein interaction mapping via MS2 or Cas13-based APEX targeting

**DOI:** 10.1101/2020.02.27.968297

**Authors:** Shuo Han, Boxuan Simen Zhao, Samuel A. Myers, Steven A. Carr, Chuan He, Alice Y. Ting

## Abstract

RNA-protein interactions underlie a wide range of cellular processes. Improved methods are needed to systematically map RNA-protein interactions in living cells in an unbiased manner. Capitalizing on the ability of the engineered peroxidase APEX2 to identify protein interaction partners via proximity-dependent biotinylation, we used two approaches to target APEX2 to specific cellular RNAs. Both an MS2-MCP system and an engineered CRISPR-Cas13 system were able to deliver APEX2 to the human telomerase RNA hTR with high specificity. One-minute proximity biotinylation captured endogenous protein interaction partners of hTR, including more than a dozen proteins not previously linked to hTR. We validated the unexpected interaction between hTR and the *N*^6^-methyladenosine (m^6^A) demethylase ALKBH5. Further investigation showed that endogenous hTR is modified by m^6^A, which can be erased by ALKBH5, and that ALKBH5 influences both telomerase complex assembly and activity. These results highlight the ability of MS2- and Cas13-targeted APEX2 to identify novel RNA-protein interactions in living cells.

## Introduction

Mapping networks of RNA-protein interactions in living cells is necessary to enable a mechanistic understanding of RNA processing, trafficking, folding, function, and degradation^1,2^. While many protein-centered approaches, such as CLIP (cross-linking immunoprecipitation)-seq and RIP (RNA immunoprecipitation)-seq, are available for the identification of RNAs bound to specific proteins of interest, few robust methods exist for the reverse problem: RNA-centered identification of protein binding partners of specific cellular RNAs of interest. Previous approaches that rely on aptamer^3–5^ or antisense probe^6–8^ affinity purification of RNA-protein complexes yield non-specific hits that bind to denatured RNAs post-lysis, and/or miss weak or transient interactions^9–11^. In addition, unbiased RNA detection is generally easier than unbiased detection of proteins due to the wide availability of RNA sequencing methods. To address the challenge of RNA-centered interactome mapping, recent studies^12,13^ have started to employ proximity-dependent labeling^14^, targeting the promiscuous biotin ligase BioID^15^ to specific MS2- or BoxB stem-loop-tagged RNAs. However, the low catalytic efficiency of BioID necessitates very long labeling times (18-24 hours), precluding the study of dynamic processes, while the stem-loop tags could affect the function of target RNAs.

Here, we present alternative methodologies for mapping RNA-protein interactions inside living cells, using both MS2 and CRISPR-Cas13^16,17^ strategies to target the proximity labeling enzyme APEX2^18^ to specific RNAs. The rapid 1-minute promiscuous biotinylation reaction catalyzed by APEX enables proteomic identification^19–23^ of endogenous interaction partners with much faster temporal resolution than BioID. We applied these methods to the human telomerase RNA (hTR), which plays a critical role in regulating cellular senescence and oncogenesis by serving as the template for reverse transcription of telomeres^24,25^. While hTR’s interaction with the telomerase complex has been extensively characterized^26^, hTR is present in stoichiometric excess over telomerase in cancer cells^27^ and is broadly expressed in tissues lacking telomerase protein^28^. These observations suggest that additional telomerase-independent hTR interactors may exist and serve to regulate hTR function^29,30^, although they have not been systematically explored. Our proximity labeling experiment recovered not only known interactors of hTR, but also novel interaction partners, including the RNA m^6^A demethylase ALKBH5^31^, highlighting the utility of RNA-targeted APEX for unbiased discovery.

## Results

### Development of an MS2-MCP strategy to target APEX to tagged hTR

We envisioned two complementary approaches to deliver APEX to the site of telomerase RNA (hTR) for proximity labeling (Figure 1A). The first entails conjugating hTR to the bacteriophage MS2 RNA stem-loop, which can specifically bind an MS2 coat protein-fused APEX2 (MCP-APEX2) with high affinity (K_d_ <1 nM)^32^. In the second approach, a catalytically inactive Cas13-APEX2 fusion (dCas13-APEX2) is programmed using a guide RNA (gRNA) to target unmodified hTR, but with lower binding affinity (K_d_ ~10 nM)^33^.

**Figure 1.**
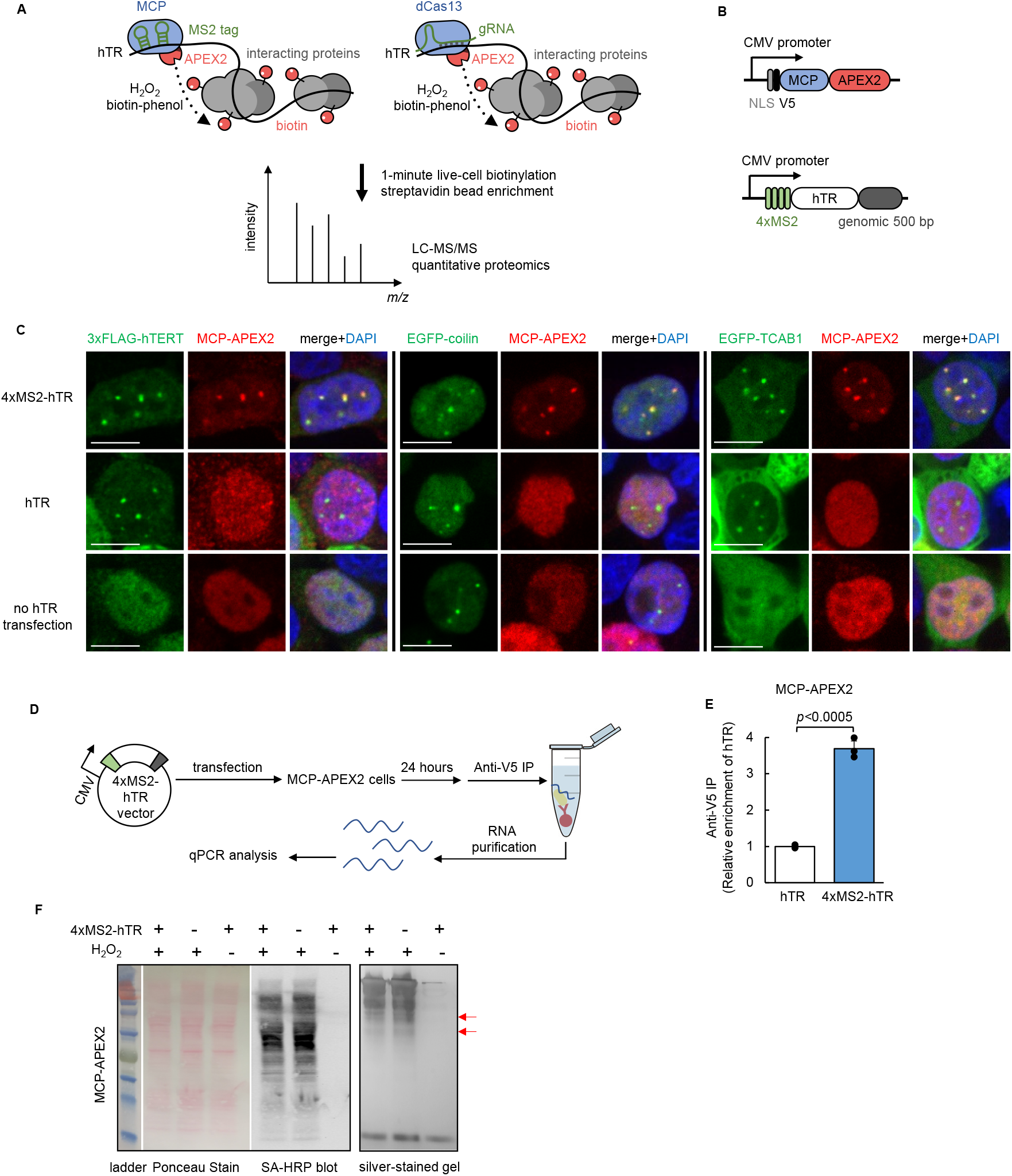
Targeting APEX2 to human telomerase RNA with MS2-MCP. **Figure 1A** Scheme of APEX2-mediated proximity labeling of human telomerase RNA (hTR) interacting proteins. Left: APEX2 is targeted to MS2 stem-loop-tagged hTR via fusion to MS2 coat protein (MCP). Right: APEX2 is targeted to hTR via fusion to catalytically dead Cas13 protein (dCas13) and a guide RNA (gRNA). To initiate labeling, H_2_O_2_ is added for 1 minute to cells preloaded with biotin-phenol, which is oxidized by APEX2 into a phenoxyl radical to covalently tag proximal endogenous proteins. Biotinylated proteins are enriched with streptavidin beads, then identified by liquid chromatography and tandem mass spectrometry (LC-MS/MS). **Figure 1B** Design of MCP-APEX2 and 4×MS2-hTR expression constructs. Genomic 500 bp is the 500 bp genomic sequence flanking the 3’ of hTR. **Figure 1C** Fluorescence imaging of MCP-APEX2 localization to hTR foci. HEK293T cells are transfected with either 4×MS2-hTR (top), untagged hTR (middle), or no hTR (bottom). MCP-APEX2 is visualized by anti-V5 staining (Alexa Fluor 647, middle column). hTR foci are visualized with either co-transfected 3×FLAG-hTERT (anti-FLAG staining, Alexa Fluor 488, left image set), co-transfected EGFP-coilin (middle image set), or co-transfected EGFP-TCAB1 (right image set). Nuclei are stained with DAPI. Scale bars, 10 μm. **Figure 1D** Scheme of RNA immunoprecipitation for validation of MCP-APEX2 targeting to hTR. HEK293T cells stably expressing MCP-APEX2 are transfected with 4×MS2-hTR or untagged hTR. RNA bound to APEX2 is immunoprecipitated with anti-V5 antibody followed by RT-qPCR quantitation. **Figure 1E** RT-qPCR of RNA immunoprecipitation following experiment in (D). The fold enrichment of hTR in cells transfected with 4×MS2-hTR is normalized against the untagged hTR control. Data are analyzed using a one-tailed Student’s t-test (n=3). **Figure 1F** Biochemical analysis of biotinylated proteins from HEK293T cells stably expressing MCP-APEX2. Lysates were run on SDS-PAGE and analyzed by Ponceau Stain (left) or streptavidin blotting (middle). Total eluted protein after streptavidin bead enrichment is visualized by silver staining (right). Red arrows point to proteins differentially biotinylated by targeted versus nontargeted APEX2.

To validate the MS2-MCP strategy, four tandem MS2 stem-loops were inserted at the 5’ end of hTR (4×MS2-hTR, Figure 1B), which is known to be flexible and single-stranded^26^, in contrast to hTR’s 3’ end. Furthermore, the 5′ end of hTR (but not 3’ end) is not processed when expressed from a Pol II promoter (e.g. CMV promoter), so that the MS2 tags remain intact in the mature transcript^34^. The mature hTR sequence is followed by 500 bp of downstream genomic sequence to ensure proper processing of the 3’ end^34^. Because fully-processed hTR accumulates in the evolutionarily-conserved subnuclear compartment Cajal body (CB)^35,36^, we used fluorescence imaging with CB markers to check for both functional hTR and recruitment of MCP-APEX2 to hTR foci. We co-expressed 4×MS2-hTR with a nuclear-localized MCP-APEX2 (Figure 1B) in HEK293T cells and performed immunostaining (Figure 1C). We found that MCP-APEX2 specifically co-localizes with the core CB component coilin when MS2 tagged hTR is co-expressed, but not in controls with untagged hTR or no hTR (Figure 1C). In addition, co-localization of MCP-APEX2 to hTR foci was confirmed by co-staining against the telomerase proteins hTERT (human telomerase reverse transcriptase) and TCAB1 (telomerase Cajal body protein 1) (Figure 1C).

As a separate readout of MCP-APEX2 targeting to 4×MS2-hTR, we performed RNA immunoprecipitation (RIP) followed by quantitative RT-PCR (Figure 1D). Consistent with the imaging data, 3.7-fold greater hTR transcripts were enriched by MCP-APEX2 (anti-V5) pull-down in cells expressing 4×MS2-hTR compared to cells expressing the untagged hTR control (Figure 1E). Together, the imaging and RIP experiments demonstrate the efficient targeting of MCP-APEX2 to hTR.

We assessed the promiscuous labeling activity of MCP-APEX2, targeted to hTR, by Western blot. Figure 1F shows H_2_O_2_-dependent biotinylation in HEK293T cells. Interestingly, biotinylated proteins enriched from cells expressing 4×MS2-hTR versus untagged hTR showed subtle differences in banding patterns, or “fingerprints”, on the silver-stained gel, consistent with the expectation that hTR-targeted APEX2 tags a somewhat different cohort of proteins than untargeted MCP-APEX2, which distributes throughout the nucleus.

To probe the generality of MS2/MCP-based APEX2 targeting, we performed imaging of a different MS2-tagged RNA, ATP5B-3’UTR-2×MS2. This RNA is known to localize to stress granules upon NaAsO_2_ treatment ^37^. Supplementary Figure 1 shows that MCP-APEX2 co-localizes with the stress granule marker G3BP1 specifically in the presence of the MS2 tagged transcript but not with the untagged control, indicating that the MS/MCP approach is also feasible for targeting APEX2 to mRNAs.

### Development of a CRISPR-Cas13 strategy to target APEX to untagged hTR

The discovery of RNA-directed CRISPR systems^33,38^ offers the exciting opportunity to target unmodified, native cellular RNAs. Thus, we next explored the use of Cas13 to deliver APEX to untagged telomerase RNA. In comparison to the small size of MCP (14 kDa), the bulk of Cas13 enzymes (~120 kDa) poses a potential steric challenge to identifying accessible sites on the highly structured hTR RNA without perturbing its endogenous interactions. Previous studies have established several Cas13 orthologs that are catalytically active inside mammalian cells, including LwaCas13a (139 kDa)^16^, PspCas13b (128 kDa)^39^, and RfxCas13d (112 kDa)^17^. We selected the smallest and most active of these, RfxCas13d^17^, for testing. We first designed a total of six gRNAs (gRNA1-6, Supplementary Figure 2) targeting different regions of hTR and validated their targeting by knockdown with an active RfxCas13d. All six gRNAs significantly reduced hTR levels in HEK293T cells, with efficiency varying from 20% to 75% (Supplementary Figure 3). However, co-expressing the gRNAs with a catalytically dead RfxCas13d (dRfxCas13d, Figure 2A) did not result in observable enrichment of dRfxCas13d at hTR foci by imaging (Figure 2B and Supplementary Figure 4). A similar test with the catalytically dead PspCas13b^39,40^ using position-matched gRNAs also failed to give enrichment at hTR foci (Supplementary Figure 5). Based on these negative results, we hypothesized that dRfxCas13d binding to hTR may require further optimization to improve its affinity.

**Figure 2.**
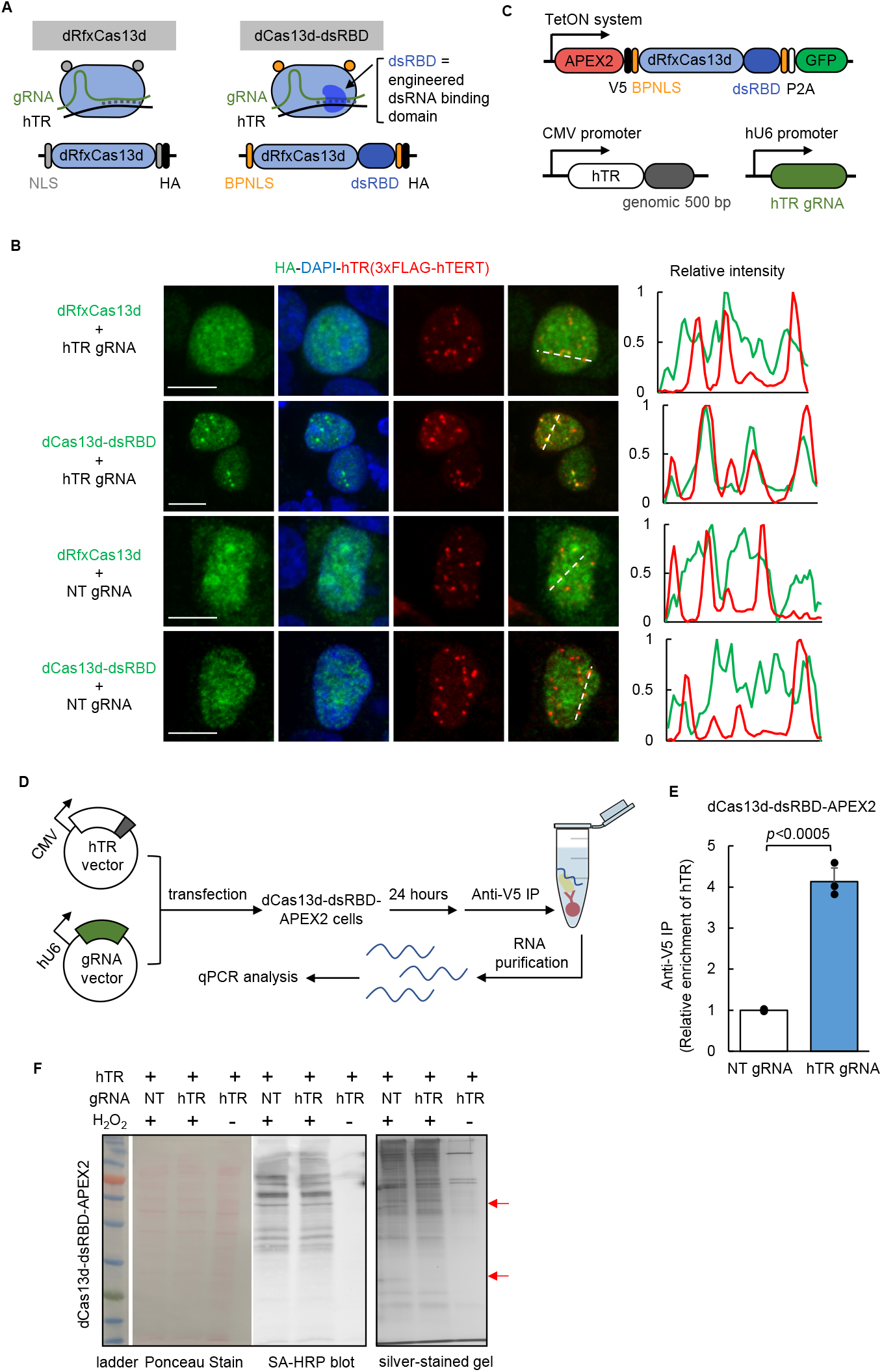
Targeting APEX2 to human telomerase RNA with CRISPR-Cas13. **Figure 2A** Design of dCas13d-dsRBD for improved hTR targeting. A sequence-independent double-stranded RNA binding domain (dsRBD) from the human protein kinase R (PKR) is fused to the C-terminus of dRfxCas13d protein to further stabilize the hTR-gRNA RNA duplex. A bipartite nuclear localization sequence (BPNLS) is used in place of the SV40 NLS for improved nuclear localization. Supplementary Figure 4 shows details of the optimization process. **Figure 2B** Fluorescence imaging of dCas13d-dsRBD localization. HEK293T cells expressing the constructs shown in (A) along with hTR marker 3xFLAG-hTERT were fixed and stained with anti-HA antibody to visualize dCas13d (Alexa Fluor 488, green), anti-FLAG antibody to visualize hTR foci (Alexa Fluor 647, red), and DAPI (nuclei). Pixel intensity plots of the dashed lines are shown at right. Scale bar, 10 μm. Supplementary Figure 6 shows additional fields of view. **Figure 2C** Design of dCas13d-dsRBD-APEX2, hTR, and gRNA expression constructs used in this study. **Figure 2D** Scheme of RNA immunoprecipitation for validation of dCas13d-dsRBD-APEX2 targeting to hTR. HEK293T cells stably expressing dCas13d-dsRBD-APEX2 are transfected with hTR and the indicated gRNA. RNA bound to APEX2 is immunoprecipitated with anti-V5 antibody followed by RT-qPCR quantitation. **Figure 2E** RT-qPCR results from the experiment in (D). The fold enrichment of hTR in cells transfected with hTR gRNA is normalized against the nontargeting (NT) gRNA control. Data are analyzed using a one-tailed Student’s t-test (n=3). **Figure 2F** Biochemical analysis of biotinylated proteins from HEK293T cells stably expressing dCas13d-dsRBD-APEX2. Lysates were run on SDS-PAGE and analyzed by Ponceau Stain (left) or streptavidin blotting (middle). Total eluted protein after streptavidin bead enrichment is visualized by silver staining (right). Red arrows point to proteins differentially biotinylated by targeted versus nontargeted APEX2.

**Figure 3.**
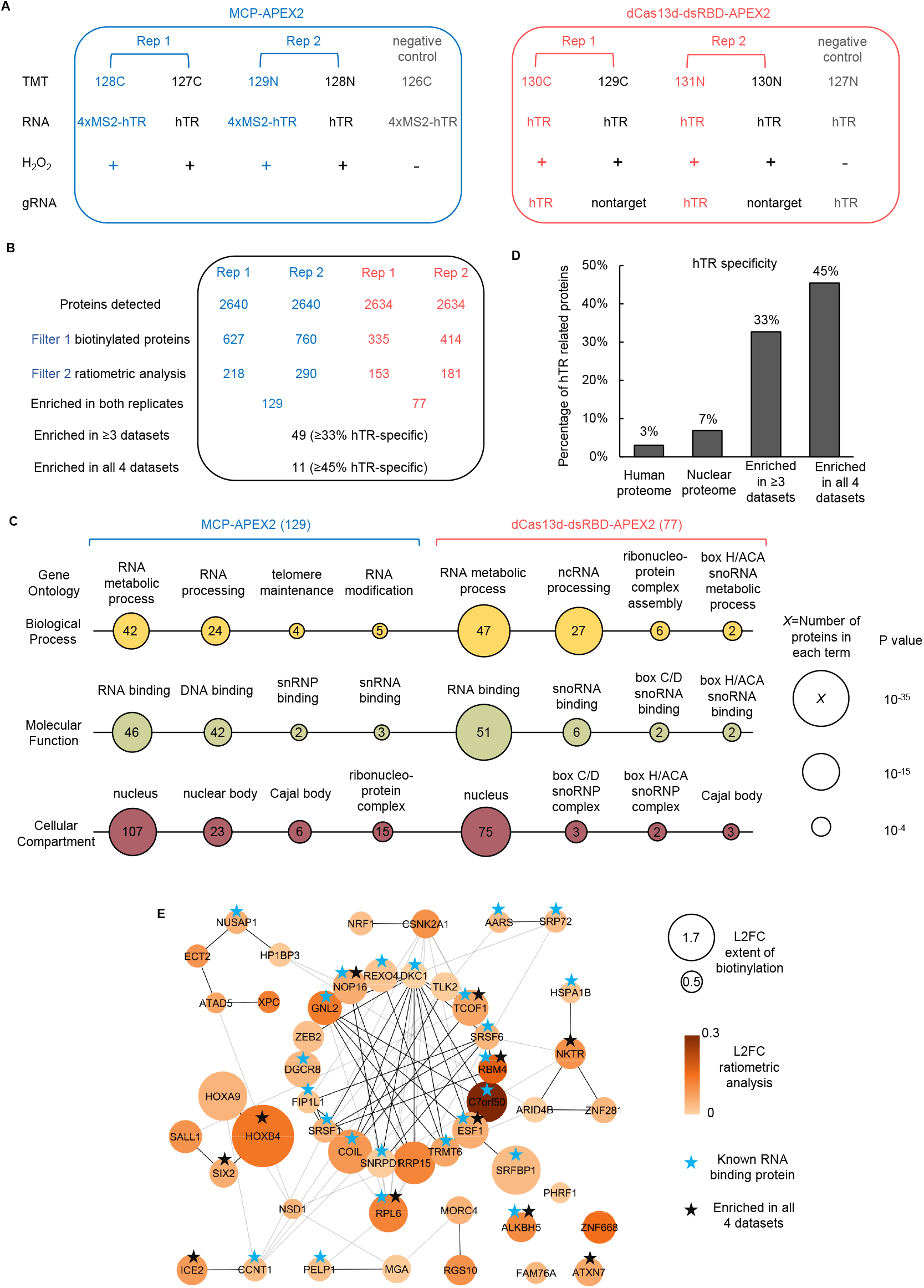
Proteomic analysis of hTR interactome via proximity labeling. **Figure 3A** Design of the proteomic experiment. HEK293T cells stably expressing the indicated APEX2 constructs are transfected with corresponding hTR and gRNA, and then labeled with biotin-phenol as in Figure 1A. Tandem mass tags (TMT) are used for quantitative proteomic analysis. Each experimental set contains two replicates and negative controls with nontargeted APEX2 or H_2_O_2_ omitted. **Figure 3B** Filtering of the proteomic data to identify enriched hTR interacting proteins. The table shows the number of proteins remaining after each filtering step. hTR specificity is calculated from interaction partners of known hTR binding proteins (list in Figure S2A) in the BioGRID database (see Methods). Scatter plots in Figure S2B show a correlation between TMT ratios across replicates. **Figure 3C** Gene ontology (GO) analysis of proteins enriched by MCP-APEX2 (left) and dCas13d-dsRBD-APEX2 (right). Only a subset of significantly enriched GO terms related to hTR is shown. Node size scales with −log_10_ (P-value). The number of proteins associated with each term is indicated inside each node. **Figure 3D** hTR specificity analysis, calculated as in (B), for the total human proteome (20996 proteins), nuclear proteome (7530 proteins), proteins enriched in ≥3 of our proteomic datasets (49 proteins), and proteins enriched in all 4 datasets (11 proteins). Specificity is defined as in Figure 2B. **Figure 3E** Protein-protein interaction (PPI) map of proteins enriched in ≥3 of our proteomic datasets. Known RNA binding proteins are marked with blue stars. Proteins enriched in all 4 proteomic datasets are marked with black stars. Node size scales with protein biotinylation extent and node color scales with the ratio of enrichment in targeting vs nontargeting control. Markov clustering was performed with PPI scores from the STRING database (see Methods).

We speculated that the interaction between dRfxCas13d and target RNA might be specifically enhanced by introducing a sequence-independent double-stranded RNA binding domain (dsRBD)^41^ to recognize the gRNA-target RNA duplex (Figure 2A) that forms only when the cognate target is bound. Such an interaction might help to cooperatively stabilize the dRfxCas13d-gRNA-target RNA ternary complex, whose structure^42^ features a solvent-exposed dsRNA backbone accessible for additional binding. To test this strategy, we selected the dsRBD from human protein kinase R (PKR), which binds indiscriminately to dsRNA ≥ 16 bp^43–45^ during viral infection, and fused it to dRfxCas13d to stabilize the 22 bp gRNA-target duplex. We found that, indeed, the addition of the dsRBD produced detectable enrichment of dRfxCas13d at nuclear foci by imaging (Supplementary Figure 4). We then optimized the site of dsRBD fusion, nuclear localization sequence, and length of the linker before arriving at the final design (dCas13d-dsRBD, Supplementary Figure 4). Remarkably, clear localization of dCas13d-dsRBD to the expected hTR foci was only observed with gRNA_5, which targets nucleotides 148-169 in the single-stranded J2a/3 region of hTR. This region is highly variable in length and sequence across vertebrates^46^, suggesting that it might play less of an important structural role than other conserved regions and serve as an accessible binding site for dCas13. In a side-by-side comparison with the starting template dRfxCas13d^17^, the optimized dCas13d-dsRBD formed nuclear foci that tightly overlapped with hTR foci marked by hTERT in the presence of gRNA_5, whereas non-targeting gRNA or original dRfxCas13d only yielded diffuse nuclear signal (Figure 2B). The greater binding capability of dCas13d-dsRBD was additionally confirmed by RNA immunoprecipitation (RIP) using another non-coding RNA, MALAT1, as target (Supplementary Figure 7).

Having established the dCas13-dsRBD based approach to target untagged hTR, we then generated an APEX2 fusion (dCas13d-dsRBD-APEX2, Figure 2C) for proximity labeling. hTR binding to dCas13d-dsRBD-APEX2 was quantified by RIP-qPCR (Figure 2D), similar to our analysis of MCP-APEX2. We found that pull-down of dCas13d-dsRBD-APEX2 with target gRNA resulted in >4-fold greater enrichment of hTR compared to a control using non-target gRNA (Figure 2E). The expected targeting of dCas13d-dsRBD-APEX2 fusion to hTR foci in the nucleus was additionally confirmed by fluorescence microscopy (Supplementary Figure 8).

We then performed a 1-minute proximity labeling reaction in HEK293T cells expressing dCas13d-dsRBD-APEX2, followed by streptavidin bead enrichment of biotinylated proteins. The streptavidin-HRP blot in Figure 2F shows that dCas13d-dsRBD-APEX2 has robust H_2_O_2_-dependent biotinylation activity. In addition, the silver-stained gel of biotinylated proteins enriched from cells co-expressing hTR gRNA versus non-target gRNA shows noticeable differences in banding pattern, in line with the expectation that APEX2 is targeted to distinct neighborhoods using the two different gRNAs.

### Proteomic mapping of hTR interacting partners via APEX proximity labeling

Previous studies have suggested that hTR may have roles beyond serving as the RNA template for telomerase. However, the hTR interactome has not been systematically examined due to the shortage of methods for RNA-centered interactome mapping. Therefore, we performed proteomic analysis of hTR-interacting proteins in living HEK293T cells, using the APEX RNA-targeting strategies developed above. We designed a 10-plex tandem mass tag (TMT)-based quantitative proteomic experiment, in which half the samples used the MS2-MCP based targeting strategy, while the other half used CRISPR-Cas13 to target APEX to hTR. Within each sample set, we performed two biological replicates of targeted APEX labeling (Figure 3A, red/blue), and two biological replicates of the negative control with untargeted APEX (Figure 3A, black). Finally, we included negative control samples (Figure 3A, grey) that omitted H_2_O_2_ in order to identify endogenously biotinylated proteins and non-specific binders to the streptavidin beads. Clonal HEK293T cells stably expressing the indicated APEX fusion constructs were incubated with the biotin-phenol probe for 30 minutes prior to the 1-minute H_2_O_2_ labeling reaction. All ten samples were independently lysed, and their biotinylated proteomes enriched using streptavidin magnetic beads. Proteins were digested on-bead to peptides with trypsin, chemically labeled with TMT reagents, pooled, and then analyzed by liquid chromatography-tandem mass spectrometry analysis (LC-MS/MS)^47^.

The proteomic experiment identified more than 2600 proteins with 2 or more unique peptides (Figure 3B). Both MCP-APEX2 and dCas13d-dsRBD-APEX2 samples exhibited a high correlation between biological replicates (Supplementary Figure 9, R^2^>0.9 for all). To remove potential contaminants and identify enriched proteins, we adopted a two-step, ratiometric filtering strategy used in our previous APEX studies^22,48,49^. Because our APEX2 constructs are localized to the nucleus, we first filtered the data by using prior nuclear annotation (possible true positive) or prior mitochondrial or endoplasmic reticulum annotation (possible false positive) as a guide (Figure 3B, Filter 1). Histograms of detected proteins plotted by 128C/126C or 130C/127N TMT ratio show preferential biotinylation of nuclear proteins over false positives (Supplementary Figure 10). We applied a false-discovery-rate (FDR) cutoff of 2% to remove potential contaminants. The remaining proteins were then filtered based on their preferential biotinylation by targeted versus untargeted APEX (Figure 3B, Filter 2). *Bona fide* hTR-interacting proteins should show greater enrichment by hTR-targeting APEX2 compared to non-targeting control (log_2_ fold change>0), as seen for known hTR-binding partners DKC1^50^ and DGCR8^51^ (Supplementary Figure 10). We further intersected the enriched proteins from the two biological replicates in each experiment to minimize contaminants and obtained final lists of 129 and 77 proteins in the MCP-APEX2 and dCas13d-dsRBD-APEX2 experiments, respectively (Figure 3B).

To separately assess the interactomes identified by MCP-APEX2 and dCas13d-dsRBD-APEX2, we first analyzed the protein lists using Gene Ontology (Figure 3C). Both lists showed high enrichment of RNA binding and RNA processing proteins in the nucleus, as expected. Notably, proteins related to the Cajal body and small nuclear/nucleolar RNA binding, which are key in regulating hTR biogenesis and processing, were also significantly enriched. These results are consistent with our fluorescence imaging and immunoprecipitation assays above. Interestingly, the analysis also revealed protein groups unique to each dataset. For example, telomere maintenance, DNA binding, and RNA modification proteins were only identified by MCP-APEX2. Such differences may be due to the use of MS2-tagged versus untagged RNA, or the different positions occupied by APEX on the hTR RNA (5’ end for MS2 versus J2a/3 region for Cas13). The high specificity of our datasets was additionally confirmed by analyzing the enrichment of known hTR binding proteins and their interaction partners^52^ (Supplementary Table 1). The hTR specificity of the 49 proteins identified in three or more of our proteomic datasets and 11 proteins identified in all four datasets is 33% and 45%, respectively (Figure 3D). In comparison, less than 3% of the human proteome and 7% of the nuclear proteome are specific to hTR using the same metrics. For the enriched proteins with no known interaction with hTR (Supplementary Table 4), literature evidence suggests that some might be connected indirectly to hTR biology. For example, RPL6, a nucleolus protein enriched in all 4 datasets, binds directly to nucleolin and NOP2^52^, both of which regulate telomerase function^53,54^. Altogether, our datasets are highly enriched for hTR-related proteins (i.e. the 33% and 45% specificities represent lower bounds) and may serve as a rich source of novel hTR-interacting protein candidates.

To further analyze the 49 proteins we enriched in three or more datasets, we performed clustering based on prior protein-protein interaction evidence in the STRING database^55^ (Figure 3E). The largest cluster contains almost exclusively RNA binding proteins (15 out of 18), including the key telomerase component DKC1, the hTR degradation complex component DGCR8, and the core Cajal body component COIL. Other smaller groups that closely associate with the major cluster also contain numerous proteins related to hTR, such as the Cajal body component ICE2. While our analyses demonstrate the high specificity achieved by the proteomic experiment, we find that our sensitivity is limited; only one (DKC1) of the six core components of telomerase (Supplementary Table 1) was identified. Of the remaining five proteins, four were not detected at all by the mass spectrometer in our experiment and were similarly undetected in previously published APEX nuclear proteomes^56–58^, suggesting that they are likely low-abundance proteins (e.g. TERT^27^) or lack surface-exposed tyrosines for APEX biotinylation. For known hTR-interacting proteins that were detected by the mass spectrometer, our ratiometric data analysis approach filtered out dual-localized proteins (e.g. PARN, EXOSC10 and TOE1^51,59–61^) that have high biotinylation over the negative control but low enrichment over the non-targeting APEX2 (Supplementary Figure 10). Such an analysis workflow increases specificity at the expense of sensitivity, a trade-off we also noted in our previous APEX studies of open subcellular compartments^22,48^.

### ALKBH5 binding to hTR regulates telomerase activity through m^6^A modification

We were intrigued by the RNA binding protein ALKBH5 (alkylated DNA repair protein alkB homolog 5), which was consistently enriched in all four proteomic datasets but has no prior connection to hTR biology nor known interaction with other hTR-related proteins that we enriched (Figure 3E). ALKBH5 was recently identified as an RNA *N*^6^-methyladenosine (m^6^A) demethylase, one of two known to date^31^. It catalyzes the removal of m^6^A from target RNAs and is implicated in the regulation of male fertility^31,62^, glioblastoma^63^, and breast cancer^64^. m^6^A is the most abundant internal modification in eukaryotic mRNA^65,66^, known to influence mRNA metabolism including export, translation, and degradation^67,68^, but its existence and regulatory functions on non-coding RNA are much less explored.

To validate our proteomic identification of ALKBH5 as an interaction partner of hTR, we performed an immunoprecipitation experiment against ALKBH5 to test its binding to endogenous hTR, as published ALKBH5 CLIP-seq data^69^ only exist for polyadenylated RNAs. RT-qPCR analysis in Figure 4A shows that immunoprecipitation of FLAG-tagged ALKBH5 enriches hTR but not a different nuclear non-coding RNA, HOTAIR. We next asked whether endogenous hTR is directly modified by m^6^A. A RIP experiment using anti-m^6^A antibody gave enrichment of hTR compared to negative controls (Figure 4B). We further tested if overexpression of ALBKH5 reduces m^6^A levels on hTR; indeed, anti-m^6^A RIP enriches less hTR RNA when ALKBH5 is overexpressed, suggesting that ALKBH5 activity can reduce the levels of m^6^A on hTR (Figure 4C). A control showed that the total level of hTR was unchanged upon ALKBH5 overexpression (Figure 4D).

**Figure 4.**
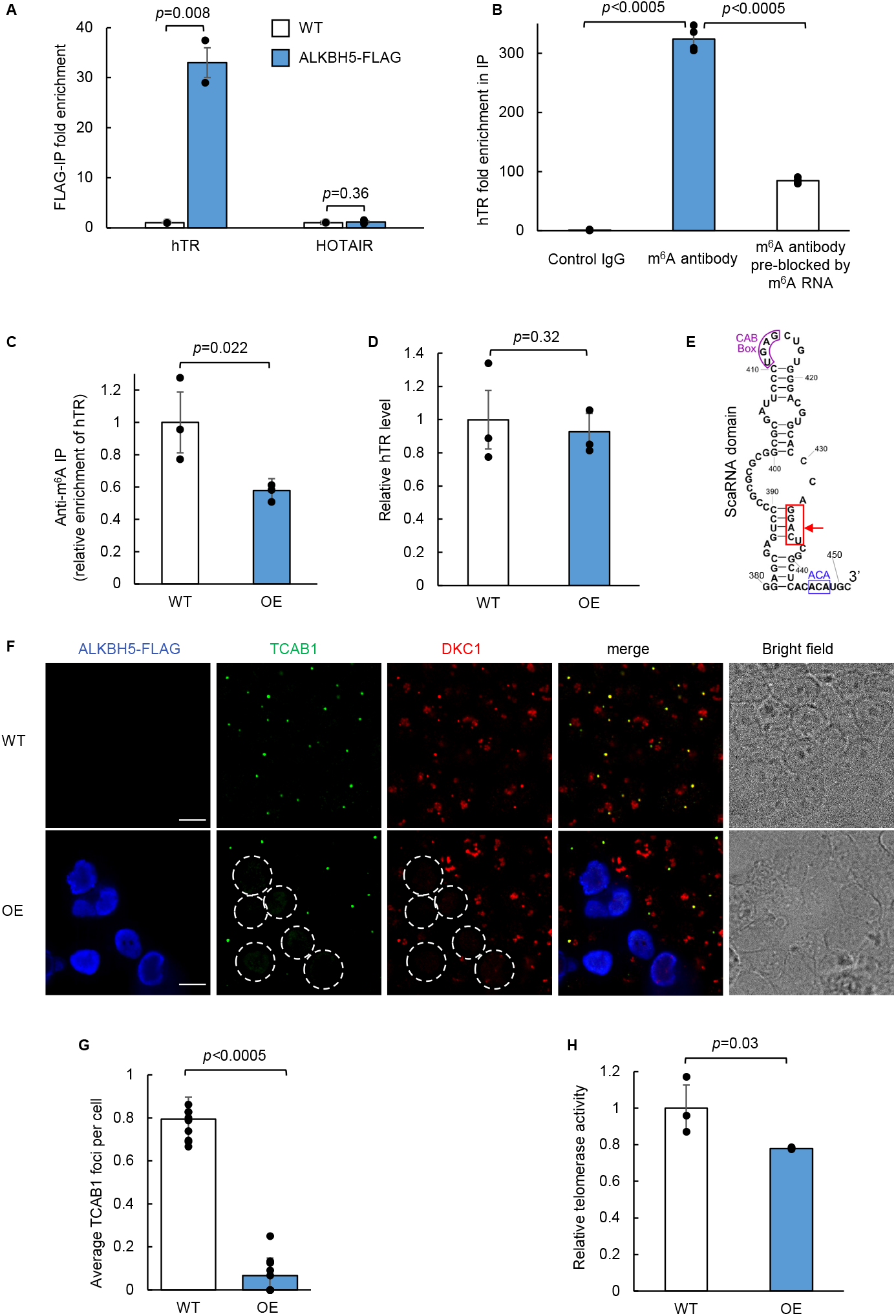
Regulation of hTR function by m^6^A demethylase ALKBH5. **Figure 4A** RT-qPCR of RNA immunoprecipitation for validation of ALKBH5 binding to hTR. Wild-type (WT) HEK293T cells are either mock-transfected or transfected with ALKBH5-FLAG. RNA bound to ALKBH5 is immunoprecipitated with anti-FLAG antibody followed by RT-qPCR quantitation. Fold enrichment of the respective RNA in ALKBH5-FLAG expressing cells is normalized against WT cells. Data are analyzed using a one-tailed Student’s t-test (n=2). **Figure 4B** RNA immunoprecipitation followed by RT-qPCR of hTR. Total RNA extract from wild-type HEK293T cells is immunoprecipitated with either a control rabbit monoclonal IgG antibody, an anti-m^6^A rabbit monoclonal IgG antibody, or the m^6^A antibody pre-blocked with a synthetic m^6^A modified luciferase RNA. The immunoprecipitated RNA is quantified by RT-qPCR. Fold enrichment of hTR is normalized against the control IgG pulldown. Data are analyzed using a one-tailed Student’s t-test (n=4). **Figure 4C** Anti-m^6^A immunoprecipitation of hTR. Total RNA extract from either wild-type (WT) or ALKBH5 overexpressing (OE) HEK293T cells is immunoprecipitated with an anti-m^6^A antibody and quantified by RT-qPCR. Fold enrichment of hTR in OE cells is normalized against the WT cells. Data are analyzed using a one-tailed Student’s t-test (n=3). **Figure 4D** RT-qPCR of hTR expression level in wild-type (WT) and ALKBH5 overexpressing (OE) cells. Total RNA extract from either WT or ALKBH5-OE HEK293T cells is quantified by RT-qPCR. hTR level in OE cells is normalized against WT cells. Data are analyzed using a one-tailed Student’s t-test (n=3). **Figure 4E** Sequence and structure of nucleotide 380-451 in hTR. The m^6^A consensus motif GGACU is highlighted in red with an arrow pointing to the putative methylated adenosine. The ACA box and CAB box motifs in the small Cajal body-specific RNA (scaRNA) domain are highlighted in blue and purple, respectively. **Figure 4F** Fluorescence imaging of telomerase component TCAB1 and DKC1 in wild-type (WT) and ALKBH5 overexpressing (OE) cells. WT or ALKBH5 OE HEK293T cells are fixed and immunostained with antibodies against endogenous TCAB1 (Alexa Fluor 488), DKC1 (Alexa Fluor 405), and ALKBH5-FLAG (anti-FLAG phycoerythrin conjugate). ALKBH5 OE cells are circled with white dotted lines. Scale bars, 10 μm. **Figure 4G** Quantitation of imaging data in (E). The total number of TCAB1 foci is divided by the total number of WT cells or by the total number of ALKBH5 OE (FLAG-positive) cells in each field of view. Data are analyzed using a one-tailed Student’s t-test (n=10). **Figure 4H** Telomerase activity assay in wild-type (WT) and ALKBH5 overexpressing (OE) cells. 1×10^6^ WT or ALKBH5 OE HEK293T cells are analyzed using a commercialized PCR-based telomerase activity assay (TRAP, Telomeric Repeat Amplification Protocol). Telomerase activity in OE cells is normalized against the WT cells. Gel showing the resulting PCR products is shown in Supplementary Figure 13. Data are analyzed using a one-tailed Student’s t-test (n=3).

If hTR is modified by m^6^A methylation, where is the modification occurring? We surveyed previous m^6^A transcriptome-wide sequencing studies (Supplementary Figure 12). Comparing datasets obtained via m^6^A antibody enrichment^70,71^ to those obtained via pull-down of various m^6^A regulators (writers METTL3 and METLL14^72^; eraser FTO^73^; nuclear m^6^A reader YTHDC1^74–76^), we found evidence for m^6^A modification in the H/ACA scaRNA domain of hTR. This domain contains a 5-nt GGACU sequence that matches the reported m^6^A consensus motif (Figure 4E). Interestingly, the adenosine within this motif (A435) is in a predicted RNA duplex region, whose secondary structure could be affected by m^6^A modification^77^. Previous studies have shown that the duplex structure of the scaRNA domain is important for the assembly of telomerase complex^26^. Hence, m^6^A modification of hTR (regulated by ALKBH5) might be a mechanism to regulate telomerase assembly and hence function.

To explore this hypothesis, we probed two telomerase protein components in the context of wild-type HEK293T cells or cells overexpressing ALKBH5 (which lowers m^6^A modification of hTR, Figure 4C). TCAB1 and dyskerin (DKC1) are both well-characterized telomerase components that directly bind hTR at the H/ACA scaRNA domain, where the postulated m^6^A modification site is located. Figure 4F shows that in wild-type HEK293T cells, endogenous TCAB1 forms discrete foci within Cajal bodies that co-localize with a subset of DKC1. However, when we overexpressed ALKBH5 in HEK293T cells, we observed a >10-fold reduction in both TCAB1 and DKC1 foci specifically in ALKBH5 overexpressing cells (outlined in Figure 4F and 4G). To rule out the possibility that ALKBH5 directly affects the mRNAs encoding telomerase components (i.e. TERT, TCAB1, NHP2, NOP10, GAR1, and DKC1), we analyzed published CLIP-seq and differentially-expressed mRNA datasets for ALKBH5^31,63,69^, and confirmed those mRNAs are not targets of ALKBH5. This suggests that ALKBH5 promotes the disassembly of telomerase components, possibly via removal of m^6^A on hTR. In addition, we performed an assay of telomerase activity using the TRAP (Telomeric Repeat Amplification Protocol) assay kit. We found that ALKBH5 overexpression caused a 22% reduction in telomerase activity compared to the control (Figure 4H), suggesting that ALKBH5 may also play a role in downregulation of telomerase activity.

## Discussion

Systematic characterization of the protein interactome of specific cellular RNAs can improve our understanding of the function and regulation of diverse RNAs inside living cells. Mass spectrometry-based proteomic identification of RNA interactomes, however, has been hampered by the shortage of methods to efficiently target and enrich specific cellular RNAs and their associated binding partners. Here, we developed two alternative methods for RNA-centered profiling based on MS2 or CRISPR-Cas13 targeting of APEX for 1-minute live-cell proximity labeling. Application of the two methods to hTR recovered a highly specific list of hTR interaction partners. We note that both of these approaches require careful screening and optimization of the MS2 stem-loop fusion site or CRISPR gRNA targeting region to find accessible binding sites that minimally impact the normal function of the target RNA.

Our strategies for targeting APEX to RNA should enable other applications as well, including electron microscopy^18^ and RNA-RNA interactome mapping. The latter could be achieved using the recently developed APEX-seq approach^78^, whereby APEX catalyzes the direct and covalent biotinylation of proximal RNAs. We anticipate a growing repertoire of RNA-centered proximity labeling tools to further expand our analysis of cellular RNAs.

An important limitation of our approach and other existing proximity labeling RNA-protein interaction mapping methods^12,13^ is that they have only been demonstrated on overexpressed, highly-abundant cellular RNAs. Based on previous studies using MCP-GFP for single-molecule imaging of MS2-tagged mRNAs^79–81^, it is likely that our MCP-APEX2 strategy should have sufficient sensitivity to be extensible to lower-abundance RNAs, albeit with an increase in number of MS2 step loops (to >20 stem loops, compared to the 4 used here). Our CRISPR-Cas13 approach, however, has much lower sensitivity, even with the engineering of the dsRBD domain. In this study, we were only able to observe APEX targeting to untagged hTR overexpressed via transfection, but not to endogenous hTR in HEK293T cells. While a small number of previous studies^16,40^, including by us, have used CRISPR-Cas13 for imaging endogenous RNAs, the targets have been highly abundant cellular RNAs, and we have found that these published systems are not as effective as our dCas13d-dsRBD for targeting APEX to hTR specifically (Supplementary Figure 5). Future improvements are needed to develop CRISPR-Cas13 systems that can broadly target endogenous, low-abundance cellular RNAs with high sensitivity.

The utility of our method is exemplified by the discovery and validation of the novel hTR interactor ALKBH5. Guided by our proteomic data, we showed that ALKBH5 directly binds to hTR and can regulate its m^6^A modification, likely in the H/ACA scaRNA domain where many telomerase components bind. Additional experiments further uncovered that changes in normal ALKBH5 expression levels disrupt telomerase complex assembly and function.

Although m^6^A is predominantly understood as regulator of mRNA stability and abundance, growing evidence suggests that it also has distinct functions with respect to non-coding RNAs. As demonstrated by recent work on the long non-coding RNA MALAT1 and its binding partner hnRNPC^77^, m^6^A can destabilize RNA duplexes to increase the accessibility of hairpin structures to RNA binding proteins. Interestingly, the putative methylated A (A435) in hTR is also within a duplex region and our data are consistent with methylation facilitating telomerase component binding, raising the interesting possibility that the mechanism underlying m^6^A regulation of hTR is similar to that of MALAT1.

## Methods

### Plasmid construction

The MCP-APEX2 construct (NLS-V5-MCP-APEX2) was created by cloning into the lentiviral vector pLX304 using Gibson assembly. The dCas13d-dsRBD-APEX2 construct (dox inducible APEX2-V5-BPNLS-dRfxCas13d-dsRBD-BPNLS-P2A-GFP) was created by cloning into an all in one piggybac, TREG/Tet-3G plasmid via Gibson assembly. The sequence of bipartite SV40NLS (BPNLS) is selected as previously described^82^. The dsRBD is taken from the double-stranded RNA binding motif (amino acid 2-81) in the protein EIF2AK2. The hTR expression plasmids were constructed by putting either MS2 stem-loop tagged or native hTR sequences under the CMV promoter, followed by 500 bp genomic sequence flanking the 3’ of hTR. Guide RNA sequences were selected and cloned into a human U6 promoter containing vector as previously described^17^. Spacer sequences of the gRNAs are listed in Supplementary Table 3. The ALKBH5-FLAG construct was cloned into the pcDNA3 vector. The following constructs were obtained from Addgene: FLAG-hTERT (#51631), EGFP-coilin (#36906), EGFP-TCAB1 (#64676), RfxCas13d (#109049), dRfxCas13d (#109050). The plasmid constructs and their features are listed in Supplementary Table 2.

### Cell culture, transfection, and clonal cell line construction

HEK293T cells from the ATCC (passages <25) were cultured in a 1:1 DMEM/MEM mixture (Cellgro) supplemented with 10% fetal bovine serum, 50 units/mL penicillin, and 50 mg/mL streptomycin at 37°C under 5% CO_2_. For fluorescence microscopy imaging experiments, cells were grown on 7×7-mm glass coverslips in 48-well plates. To improve the adherence of HEK293T cells, we pre-treated glass slides with 50 mg/mL fibronectin (Millipore) for 20 minutes at 37°C before cell plating and washing three times with Dulbecco’s PBS (DPBS) (pH 7.4). All constructs were transfected using lipofectamine 2000 (Thermo) according to the manufacturer’s protocol. The amount of plasmids used for each experiment is detailed below. To generate the MCP-APEX2 clonal cells, a stable population was first created by infecting HEK293T cells with lentivirus containing MCP-APEX2 followed by blasticidin selection (8 μg/ml). Single-colonies were then picked from the stable population using glass cloning cylinders (Millipore). To generate the dCas13d-dsRBD-APEX2 clonal cells, a stable population was first created by transfecting HEK293T cells with the dCas13d-dsRBD-APEX2 plasmid, followed by puromycin selection (1 μg/ml). Single-colonies were then picked from the stable population based on their GFP fluorescence upon addition of doxycycline (400 ng/ml).

### APEX labeling

HEK293T cells stably expressing the corresponding APEX2 fusion are plated in T150 flasks at 70-80% confluency 18-24 hours prior to transfection. For the MCP-APEX2 samples, cells were transfected with 48 μg of the 4×MS2-hTR plasmid for 24 hours prior to APEX labeling. For the dCas13d-dsRBD-APEX2 samples, cells were transfected with 48 μg of the hTR plasmid and 48 μg of the guide RNA plasmid, and doxycycline (400 ng/ml) was also added to induce expression for 24 hours prior to APEX labeling. Biotin phenol (Iris Biotech) was dissolved in DMSO as 500 mM stock solution and added directly to cell culture media to a final concentration of 500 μM. After incubating the cells for 30 minutes at 37°C, H_2_O_2_ was spiked into the cell culture media to a final concentration of 1 mM to induce biotinylation. After 1 minutes of very gently swirling, the media was decanted as quickly as possible and the cells were washed three times with PBS containing 100 mM sodium azide, 100 mM sodium ascorbate and 50 mM TROLOX (6-hydroxy-2,5,7,8-tetramethylchroman-2-carboxylic acid). Cells were scraped and transferred to 15 ml Falcon tubes with ice cold PBS, spun at 3000×*g* for 5 minutes, flash frozen in liquid nitrogen and stored at −80°C.

### Streptavidin bead enrichment of biotinylated proteins

APEX labeled cell pellets from a T150 flask was used as input for each sample. The pellets were then lysed in RIPA lysis buffer (50 mM Tris, 150 mM NaCl, 0.1% SDS, 0.5% sodium deoxycholate, 1% Triton X-100, protease cocktail [Sigma-Aldrich], and 1 mM PMSF) for 5 minutes at 4°C. The lysates were cleared by centrifugation at 15,000×*g* for 10 minutes at 4C. Streptavidin-coated magnetic beads (Pierce) were washed twice with RIPA buffer, and 8 mg of each sample was separately incubated with 450 μL of magnetic bead slurry with rotation for 1 hour at room temperature or overnight at 4°C. The beads were subsequently washed twice with 1 mL of RIPA lysis buffer, once with 1 mL of 1 M KCl, once with 1 mL of 0.1 M Na_2_CO_3_, once with 1 mL of 2 M urea in 10 mM Tris-HCl (pH 8.0), and twice with 1 mL RIPA lysis buffer. For streptavidin blot analysis, biotinylated proteins were then eluted by boiling the beads in 75 μL of 3× protein loading buffer supplemented with 20 mM DTT and 2 mM biotin, and run on SDS-PAGE gel.

### Gels and western blots

Clarified lysates were boiled for 10 minutes at 95oC before loading onto an SDS-PAGE gel (Invitrogen). Gels were transferred to nitrocellulose membrane, stained by Ponceau S (10 minutes in 0.1% [w/v] Ponceau S in 5% acetic acid/water). The blots were then blocked 3% BSA in TBS-T (Tris-buffered saline, 0.1% Tween 20) at room temperature for 1 hour and stained with 0.3 μg/mL streptavidin-HRP (BioRad) in TBS-T for 1 hour at room temperature. The blots were washed four times with TBS-T for 5 min each prior to developing with Clarity Western ECL Blotting Substrates (BioRad) and imaging on a UVP BioSpectrum Imaging System. Silver-stained gels of enriched material eluted from the streptavidin beads were generated using Pierce Silver Stain Kit (Thermo).

### RNA immunoprecipitation followed by quantitative PCR

RNA immunoprecipitation experiments to test the binding between MCP-APEX2 or dCas13d-dsRBD-APEX2 and the target hTR were performed as previously described^16,83^ with minor modifications. Briefly, HEK293T cells stably expressing the corresponding APEX2 fusion are plated in 6-wells at 70-80% confluency 18-24 hours prior to transfection. For the MCP-APEX2 samples, cells were transfected with 3 μg of the 4×MS2-hTR plasmid. For the dCas13d-dsRBD-APEX2 samples, cells were transfected with 3 μg of the hTR plasmid and 3 μg of the guide RNA plasmid, and doxycycline (400 ng/ml) was also added to induce expression. 48 hours post transfection, cells were fixed with 0.1% paraformaldehyde (Ricca Chemical) in DPBS for 10 minutes at room temperature, and then quenched with 125 mM glycine in PBS for 5 minutes. Cells were lysed in RIPA buffer supplemented with protease inhibitor cocktail (Sigma-Aldrich) and RiboLock RNase inhibitor (Thermo) before centrifugation at 16,000 *g* for 10 minutes at 4°C. After saving 5% of the supernatant as input, the rest of the clarified lysate was then incubated with 25 uL Protein G magnetic beads (Pierce) coupled with 1 μL mouse anti-V5 (Life Technologies) antibody for 2 hours at 4°C with rotation. The beads were then pelleted and washed four times with RIPA buffer supplemented with 0.02% Tween-20 and RiboLock. Enriched RNAs were released from the beads in 100 μL of Elution Buffer (2% N-lauryl sarcoside, 10 mM EDTA, 5 mM DTT, in PBS supplemented with 200 μg proteinase K [Thermo] and RiboLock) at 42°C for 1 hour, followed by 55°C for 1 hour. Eluted samples were cleaned up using Agencourt RNAClean XP magnetic beads (Beckman Coulter) and eluted into 85 μL H_2_O. Thereafter, contaminating DNA was removed by digestion with RNase-free DNase I (Promega). Purified RNAs were again cleaned up using Agencourt RNAClean XP beads as above and eluted into 30 μL H_2_O. The resulting enriched RNA and input RNA is used as the template for cDNA synthesis using SuperScript III (Invitrogen) and quantified by qRT-PCR (BioRad) with Maxima SybrGreen qPCR Master Mix (Thermo) and specific primers (Supplementary Table 3). The recovery of specific RNAs is calculated by dividing transcript abundance in the enriched sample by its input.

### On-bead digestion and liquid chromatography mass spectrometry

Mass spectrometry-based proteomic experiment was performed as previously described with minor modifications^57^. Briefly, after enrichment and washing, beads were resuspended in 50 mM TRIS pH 8, 150 mM NaCl, and frozen until digestion. After thawing, dry urea was added to the solution to 8M, along with TCEP and iodoacetamide to 5 mM, followed by incubation for 30 minutes at 25°C. The beads were diluted to 4 M urea with resuspension buffer above, and 1 uL of LysC (Wako) was added and incubated at 37°C for two hours. The urea was again diluted to 2 M and 800 ng trypsin was added for overnight digestion at 37°C. The following day the digests were acidified, desalted, TMT (tandem mass tag)-labeled on-column, and fractionated according to previous reports^57^, with the exception that 100 mM potassium phosphate pH 8.0 was used in place of HEPES during the on-column TMT labeling. Six basic reverse-phased fractions were generated, concatenated to three and analyzed using an EASY-nLC 1200 coupled to a Q-Exactive Plus mass spectrometer (Thermo).

### Proteomic data analysis

LC-MS/MS data was analyzed using Spectrum Mill (Agilent) against a human Uniprot database (12/28/2017) that included 264 common laboratory contaminants. Individual TMT ratios for human proteins identified with at least two peptides were global median normalized prior to the filtering and thresholding described below. Each of the replicates was analyzed independently. To select cutoffs for proteins biotinylated by APEX2 over nonspecific bead binders (Filter 1), we first classified 3 classes of proteins according to their Gene Ontology cellular component (GOCC) annotation (10/02/2019): (1) mitochondrial matrix; (2) endoplasmic reticulum; (3) nucleus. Proteins that are annotated to be in either class (1) or (2) but not in class (3) were defined as false-positive hits, as they should not be biotinylated by our nucleus localized APEX2. We then ranked the data in each replicate by the enrichment ratio over the unlabeled negative control samples (i.e. 128C/126C, 129N/126C, 130C/127N, 131N/127N), and calculated the false-positive rate we would obtain if we retained only proteins above that TMT ratio. A false-positive rate of <2% was chosen as cutoff for Filter 1, which corresponds to cutoff at 0.4375, 0.3375, 0.6985, 0.622, for ratios 128C/126C, 129N/126C, 130C/127N, 131N/127N, respectively. To select cutoffs for proteins that were preferentially biotinylated by APEX2 targeted to hTR versus general nucleus-targeted APEX2 (Filter 2), we ranked the data in each replicate by the enrichment ratio over nucleus-targeted APEX2 samples (i.e. 128C/127C, 129N/128N, 130C/129C, 131N/130N) after the application of Filter 1. The enrichment ratio is then normalized by the median of the remaining class (3) proteins to account for differences in total protein amount between samples within the TMT experiment. Proteins with enrichment ratio log_2_ fold-change >0 were retained for further analysis. After applying Filter 1 and Filter 2 cutoffs, each of the two replicates in the MCP-APEX2 experiment or dCas13d-dsRBD-APEX2 experiment were intersected to produce the final proteomes of 129 and 77 proteins, respectively. To assess the specificity of the enriched proteins, we analyzed the hTR specificity and Gene Ontology enrichment, respectively. We defined hTR-specific proteins using a list of ten known hTR-binding proteins curated from literature (Supplementary Table 1), their direct interaction partners in the BioGrid database (https://thebiogrid.org/), and components of Cajal body according to GOCC (10/02/2019). For Gene Ontology analysis, we uploaded the final proteomes to the Gene Ontology database search portal (http://geneontology.org/, 10/02/2019) on Cellular Compartment, Biological Process, or Molecular Function to retrieve the plotted terms with their corresponding *p* values (Fisher’s Exact test with no correction). For the protein network analysis, the enriched proteins were clustered by their reported protein-protein interactions and corresponding confidence scores (STRING database, 10/02/2019) using a Markov clustering algorithm (cutoff 0.25, inflation value set at 3) and plotted in Cytoscape (v3.7.1).

### Immunofluorescence staining

HEK 293T cells were plated onto glass coverslips in 48-wells 18-24 hours prior to transfection using lipofectamine 2000. The following amount of each plasmid was used: 50-70 ng of the MCP-APEX2 plasmid, 50-70 ng of the FLAG-hTERT plasmid, 50-70 ng of the EGFP-TCAB1 plasmid, 50-70 ng of the dRfxCas13d plasmid, 50-70 ng of the dCas13d-dsRBD plasmid, 400-500 ng of the 4×MS2-hTR plasmid, 400-500 ng of the hTR plasmid, 400-500 ng of the gRNA plasmid, 300 ng of the ALKBH5-FLAG plasmid. 24 hours after transfection, the cells were fixed with 4% (v/v) paraformaldehyde in PBS buffer at room temperature for 15 minutes. Cells were then washed three times with PBS and permeabilized with cold methanol at −20°C for 5-10 min. Cells were then washed three times with PBS, and then incubated with primary antibody in PBS supplemented with 3% (w/v) BSA for 1 hour at room temperature. After washing three times with PBS, cells were then incubated with DAPI and secondary antibody in PBS supplemented with 3% (w/v) BSA for 30 minutes at room temperature. Cells were then washed three times with PBS and imaged by confocal fluorescence microscopy. The following primary and secondary antibodies were used: mouse anti-V5 (Life Technologies, 1:1000 dilution), mouse anti-FLAG (Agilent, 1:800 dilution), rabbit anti-V5 (GenScript, 1:500 dilution), rabbit anti-HA (Cell Signaling, 1:1000 dilution), FLAG-PE (Miltenyi Biotec, 1:500 dilution), rabbit anti-TCAB1 (Bethyl Laboratories, 1:500 dilution), mouse anti-DKC1 (Santa Cruz Biotech, 1:500 dilution), goat anti-mouse Alexa Fluor405 (Invitrogen, 1:1000 dilution), goat anti-mouse Alexa Fluor488 (Invitrogen, 1:1000 dilution), goat anti-mouse Alexa Fluor647 (Invitrogen, 1:1000 dilution), goat anti-rabbit Alexa Fluor405 (Invitrogen, 1:1000 dilution), goat anti-rabbit Alexa Fluor488 (Invitrogen, 1:1000 dilution), and goat anti-rabbit Alexa Fluor647 (Invitrogen, 1:1000 dilution).

### Confocal fluorescence imaging

Confocal imaging was performed using a Zeiss AxioObserver.Z1 microscope, outfitted with a Yokogawa spinning disk confocal head, a Cascade II:512 camera, a Quad-band notch dichroic mirror (405/488/568/647), 405 (diode), 491 (DPSS), 561 (DPSS), and 640 (diode) nm lasers (all 50 mW). DAPI (405 laser excitation, 445/40 emission), Alexa Fluor568 (561 laser excitation, 617/73 emission), phycoerythrin (561 laser excitation, 617/73 emission), and Alexa Fluor647 (640 laser excitation, 700/75 emission), and differential interference contrast (DIC) images were acquired through a 63× oil-immersive objective. All images were collected, processed, and analyzed using the SlideBook 6.0 software (Intelligent Imaging Innovations).

### ALKBH5 pulldown assay

One 10-cm dish of HEK293T cells were transfected with 8 μg of ALKBH5-FLAG plasmid at 70-80% confluency for 24 hours. The cells were rinsed with PBS for three times, scraped off the plate in 10 mL of PBS, and collected by centrifugation. The cell pellets were lysed with 1 mL lysis buffer (50 mM Tris-HCl pH 7.4, 150 mM NaCl, 1 mM EDTA, 1 mM dithiothreitol, 0.5% NP-40, 1:200 SUPERase• In [Invitrogen], 1:100 Protease Inhibitor Cocktails [Sigma-Aldrich]) on ice for 10 minutes, then flash frozen with liquid nitrogen and store at −80 °C. After thawing, the lysate was centrifuged at max speed and the supernatant was taken for pulldown. For target RNA pulldown, 20 μL suspension of Anti-FLAG M2 magnetic beads (Sigma-Aldrich) were washed with lysis buffer for 3 times and mixed with 500 μL of supernatant lysate. The beads mixture was incubated at 4 °C for 2 hours with rotation, washed 5 times with lysis buffer, and eluted by adding TRIzol directly. The enriched RNA and input total RNA from 5% of lysate were purified using Direct-zol RNA Purification Kit (Zymo). Target genes in input total RNA and enriched RNA were quantified by qRT-PCR using SuperScript III Platinum One-Step qRT-PCR Kit (Invitrogen). Control group with untransfected HEK cells was processed with the same protocol in parallel.

### m^6^A pulldown assay

HEK293T cells are plated in 10-cm dishes at 70-80% confluency. 18 hours after plating, cells were either directly harvested or transfected with 20 μg of the ALKBH5 plasmid with lipofectamine and then harvested 48 hours post transfection. To harvest the cells, each 10-cm dish of cells were lifted, washed with PBS for two times, and collected by centrifuge. 5mL Trizol (Thermo) was used to homogenize the cell pellets and total RNA was extracted according to manufacturer’s protocol by precipitation. m^6^A pulldown was performed with a rabbit monoclonal anti-m^6^A antibody in the EpiMark *N*^6^-Methyladenosine Enrichment Kit (NEB) according to the manufacturer’s protocol. Briefly, 80 μg of total RNA was subject to m^6^A enrichment by incubating with 25 μL Protein G magnetic beads (Pierce) coupled with 2 μL m^6^A antibody at 4 °C for 1 hour with rotation, washed 6 times, and eluted with Buffer RLT (Qiagen). Two control samples were processed in parallel: one control used 10 μL m^6^A-modified RNA (100 nM) supplied in the kit for pre-incubation with the m^6^A antibody at 4 °C for 30 minutes to block the antibody; the other used 2 μL rabbit monoclonal anti-HA antibody (Cell Signaling) as a control IgG for pulldown. The eluate was further purified and concentrated with RNA Clean and Concentrator kit (Zymo). Target genes in input total RNA and enriched RNA were quantified by qRT-PCR using SuperScript III Platinum One-Step qRT-PCR Kit (Invitrogen).

### Telomerase activity assay

HEK293T cells are plated in 10-cm dishes at 70-80% confluency. 18 hours after plating, cells were transfected with 20 μg of the ALKBH5 plasmid with lipofectamine and then harvested 48 hours post transfection. Telomerase activity is measured with the TRAPeze Telomerase Detection Kit (Millipore) according to manufacturer’s protocol. Briefly, HEK cells were lifted from plates and washed with PBS for two times, counted using Countess II FL Automated Cell Counter (Thermo) and aliquoted in 1 million cells per tube and collected by centrifugation. The cell pellets were flash frozen with liquid nitrogen and stored at −80 °C until ready for use. The cell pellets were later lysed using buffer supplied with the kit on ice for 30 minutes, centrifuged, and the supernatant was collected and aliquoted for storage in −80 °C. Each aliquot was used only once. For the telomerase activity assay, each reaction was set up in half scale (25 μL/reaction) according to the kit protocol, subjected to 32 cycles of PCR, and run on 10% TBE gel in 0.5x TBE buffer for 1 hour at 160V. The gel was then stained in 100 mL water with 10,000x SybrSafe (Thermo) stain at room temperature for 30 minutes, washed with water for 20 minutes, and imaged on a ChemiDoc MP Imaging System (Bio-rad). The DNA products were quantified with ImageJ with appropriate background correction according to the instructions in the kit.

### Bioinformatic analysis of published sequencing datasets

Reanalysis of the published datasets in Supplementary Figure 12 was carried out with IGV software. PA-m^6^A-seq and miCLIP peak data (bed files) were downloaded directly from GEO (accession number GSE54921 and GSE63753). The remaining CLIP-seq peak data were acquired using the ENCORI starBase platform^84^ (http://starbase.sysu.edu.cn/download.php). All entries were loaded into the same IGV session and aligned to hg19 reference genome. The proposed modified region at the 3’ end of hTR was shown in the figure with center motif highlighted.

## Acknowledgements

We thank Omar Abudayyeh and Jonathan Gootenberg (MIT) for help and advice on Cas13-related experiments, and members of the Ting laboratory for technical support and advice. S.H. is supported by the Stanford Bio-X Bowes Graduate Fellowship. B.S.Z was a Wu-Tsai Neurosciences Institute interdisciplinary scholar. A.Y.T is an investigator of the Chan Zuckerberg Biohub. C.H. is an investigator of the Howard Hughes Medical Institute. This work was supported by the National Institutes of Health (U24-CA210986 to S.A.C., U01-CA214125 to S.A.C., R01-DK121409 to S.A.C. and A.Y.T.) and the Wu-Tsai Neurosciences Institute of Stanford.

## Author information

### Contributions

S.H. and A.Y.T conceived this project. S.A.M performed mass spectrometry analysis of the proteomic samples and carried out initial data analysis under the advisement of S.A.C. B.S.Z performed the experiments related to ALKBH5 with input from C.H and assistance from S.H. S.H. designed and performed all the other experiments with input from A.Y.T and assistance from B.S.Z and S.A.M. S.H., B.S.Z, and A.Y.T wrote the manuscript with input from all authors.

### Competing interests

The authors declare no competing financial interests.

Correspondence and requests for materials should be addressed to (A.Y.T.).

---

**Supplementary Figure 1.**
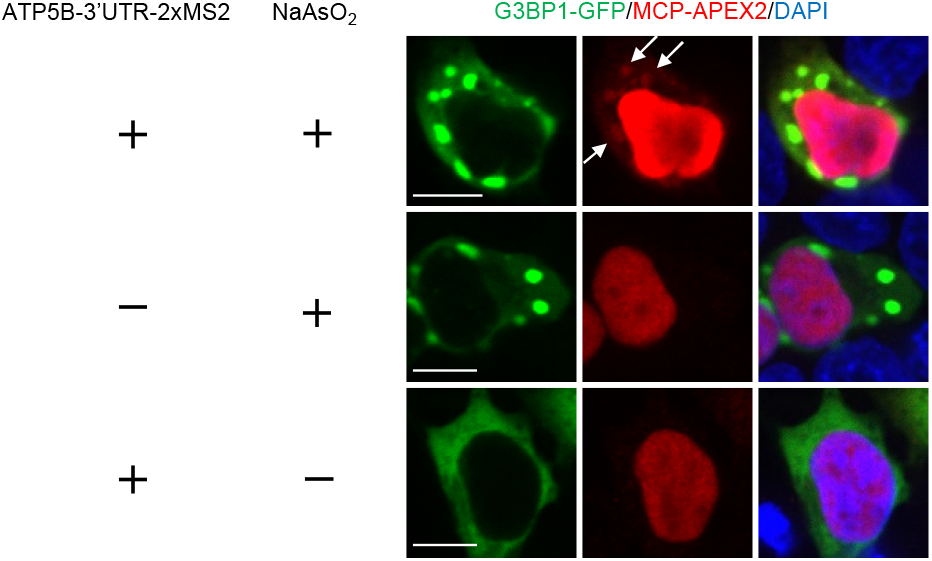
Fluorescence imaging of MCP-APEX2 binding to ATP5B transcript during stress granule formation. HEK293T cells are transfected with ATP5B-3’UTR-2xMS2 and treated with NaAsO_2_ to induce stress granule formation. MCP-APEX2 is visualized by anti-V5 staining (Alexa Fluor 647, red). Stress granules are visualized with G3BP1-GFP (green). The nucleus is stained with DAPI. White arrows show co-localization of stress granules and MCP-APEX2 only in the presence of both ATP5B-3’UTR-2xMS2 and NaAsO_2_ treatment. Scale bar, 10 μm.

**Supplementary Figure 2.**
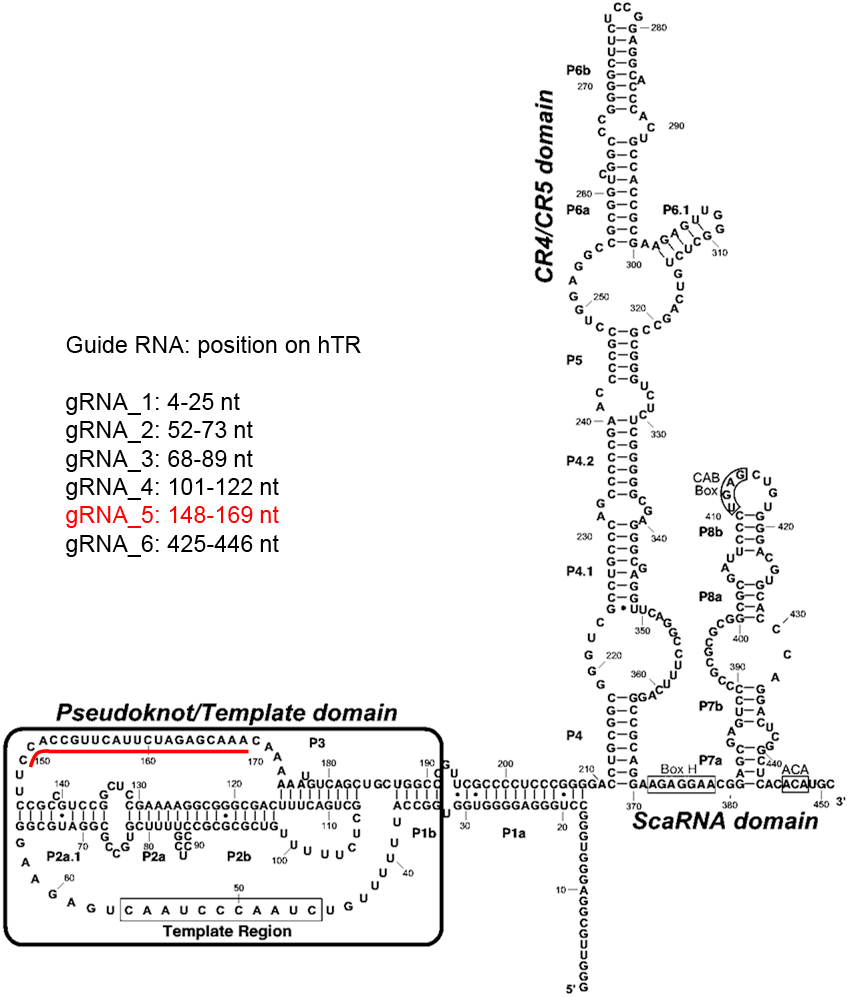
Sequence and structure of the human telomerase RNA. The spacer sequences of the RfxCas13d guide RNAs that showed significant knockdown are listed according to their position on hTR. gRNA_5 and its targeting region are highlighted in red. The structure of hTR is adapted from Chen et al Cell 2000 and http://telomerase.asu.edu/structures.html.

**Supplementary Figure 3.**
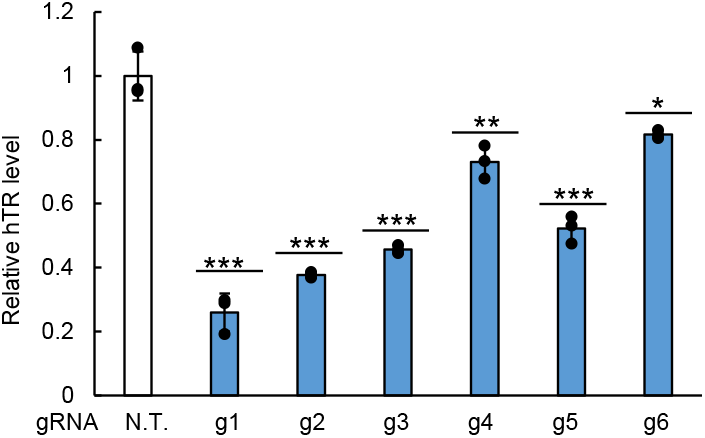
Testing hTR gRNA by knockdown with RfxCas13d. HEK 293T cells were co-transfected with RfxCas13d and the indicated gRNA. The expression level of hTR was quantified by RT-qPCR following total RNA extraction and normalized against the nontargeting (NT) gRNA control. Data are analyzed using a one-tailed Student’s t-test (n=3). *, p<0.05; **, p<0.005; ***, p<0.0005.

**Supplementary Figure 4.**
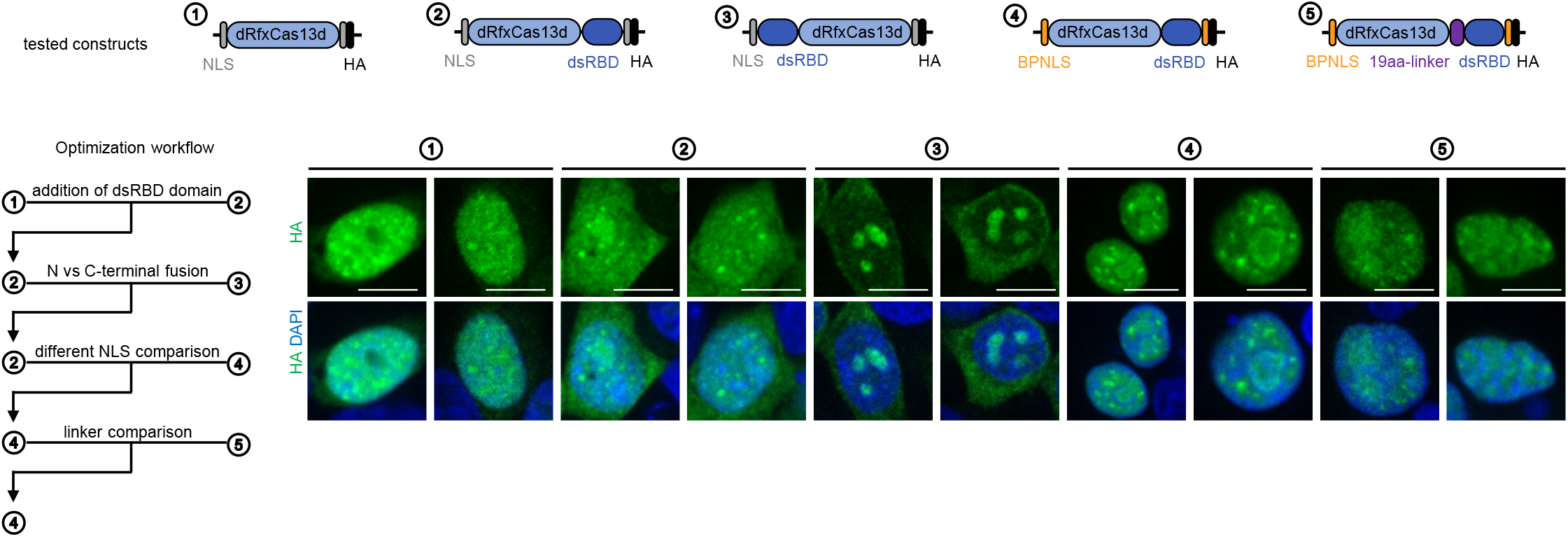
dCas13d-dsRBD optimization process. The addition of dsRBD domain to the C-terminus of dRfxCas13d improves the targeting to hTR foci: ② is better than ①. Compared to C-terminal dsRBD fusion, fusing dsRBD to the N-terminus of dRfxCas13d results in nucleolar aggregates: ② is better than ③. Replacing SV40 NLS with BPNLS improves dCas13d-dsRBD localization into the nucleus: ④ is better than ②. The addition of a flexible 19-amino acid glycine-serine linker between dCas13d and dsRBD diminishes the improved targeting: ④ is better than ⑤. All constructs are tested by co-transfection with hTR and hTR targeting gRNA in HEK293T cells. dCas13d is visualized by anti-HA staining (Alexa Fluor 488, green). The nucleus is stained with DAPI (blue). Scale bars, 10 μm.

**Supplementary Figure 5.**
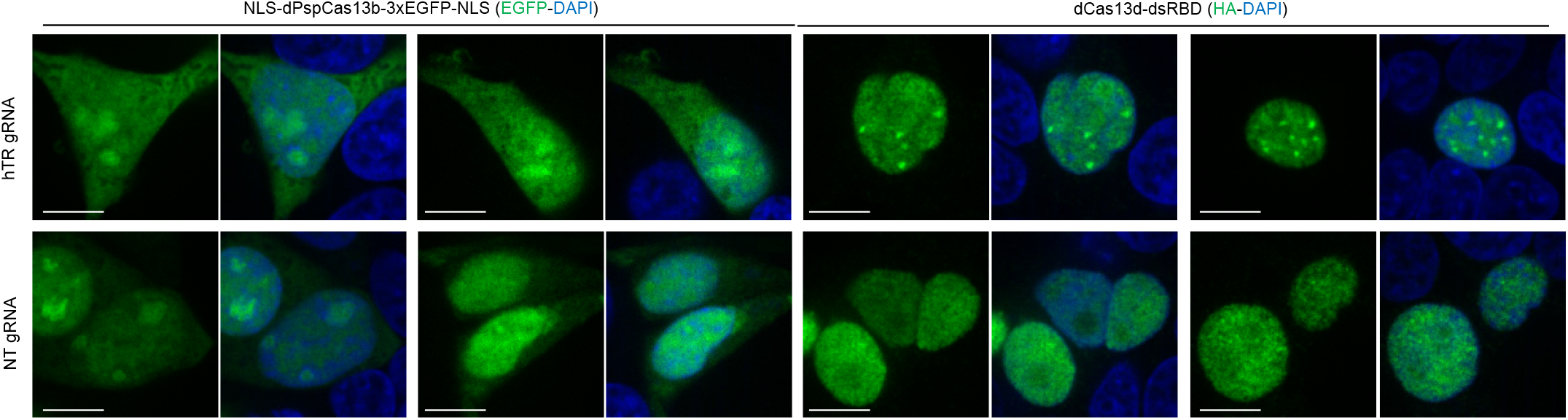
Fluorescence imaging of dPspCas13b and dCas13d-dsRBD localization to hTR. HEK293T cells were transfected with hTR, either NLS-dPspCas13b-3xEGFP-NLS or dCas13d-dsRBD, and position-matched guide RNAs. The cells were then fixed and directly imaged by EGFP (green) or stained with anti-HA antibody (Alexa Fluor 488, green) to visualize the Cas13 protein and DAPI (blue, nuclei). Scale bar, 10 μm.

**Supplementary Figure 6.**
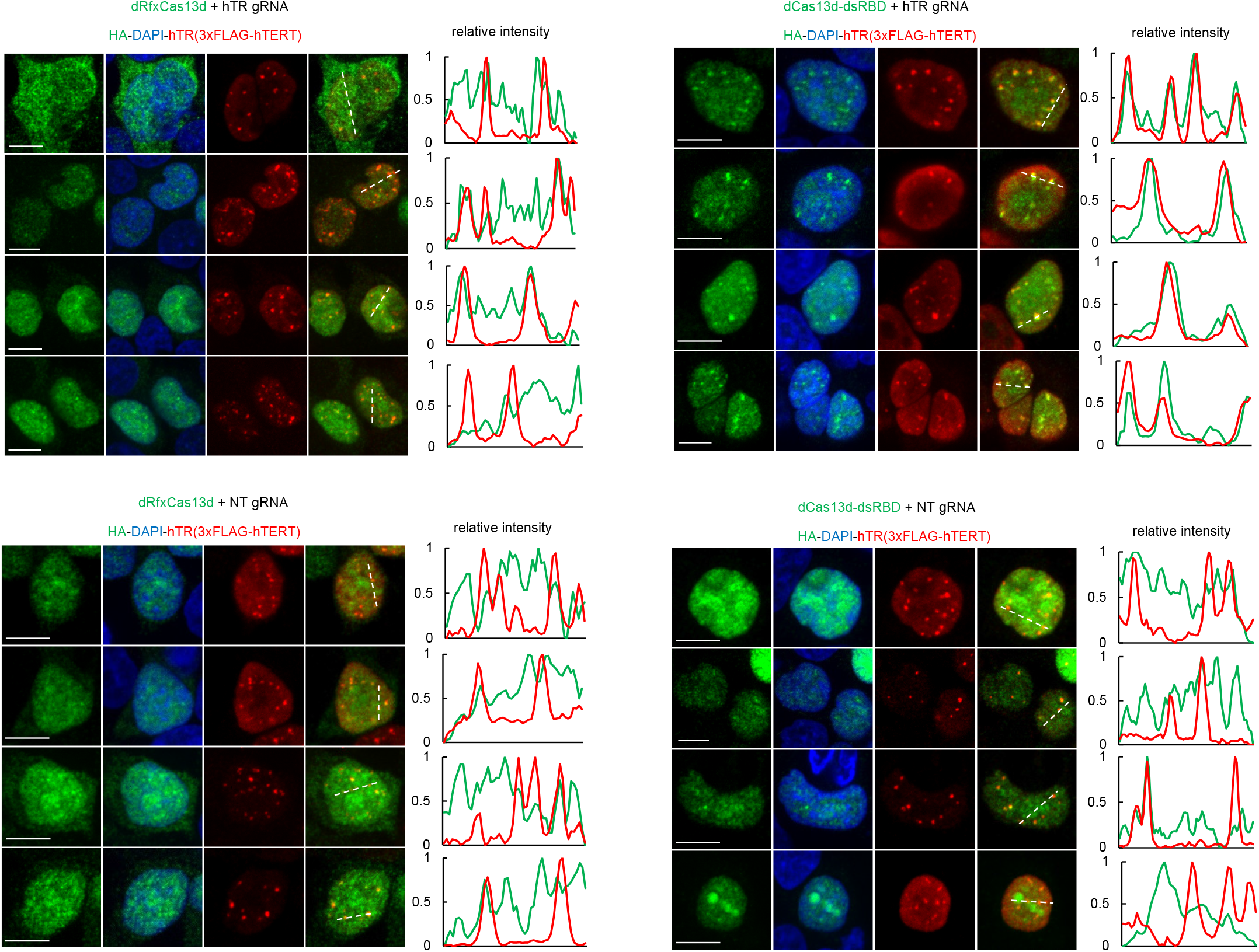
Fluorescence imaging of dCas13d-dsRBD localization. Same as in Figure 2B with additional fields of view.

**Supplementary Figure 7.**
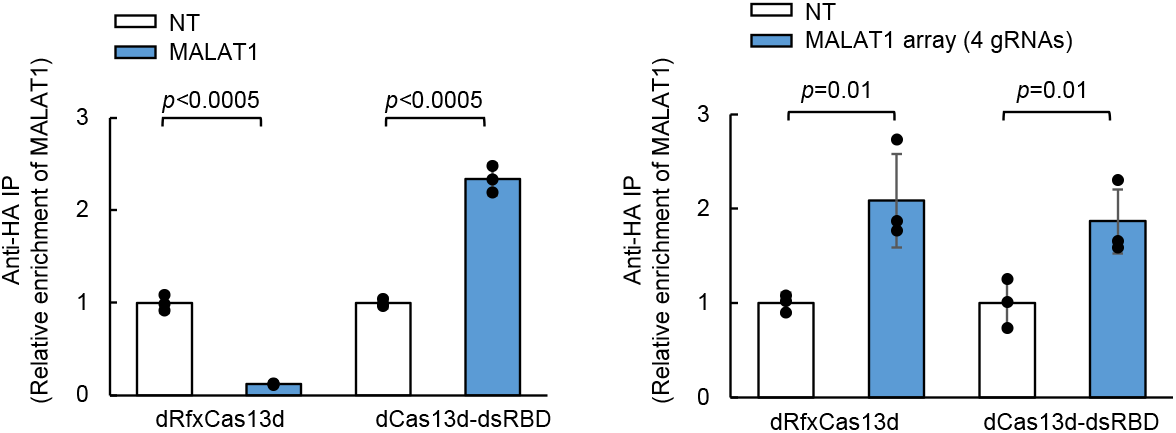
RNA immunoprecipitation experiment to show dCas13d-dsRBD binding to endogenous MALAT1 lncRNA. HEK293T cells are transfected with dRfxCas13d or dCas13d-dsRBD, together with a single MALAT1 gRNA (left) or a 4 MALAT1 gRNA array (right). RNA bound to dCas13d is immunoprecipitated with anti-HA antibody followed by RT-qPCR quantitation. Data are analyzed using a one-tailed Student’s t-test (n=3).

**Supplementary Figure 8.**
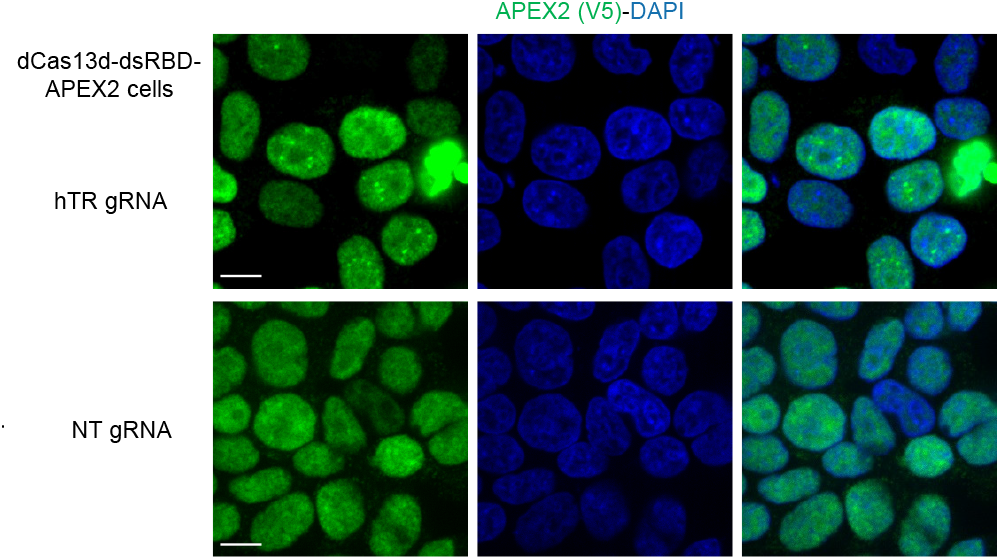
Fluorescence imaging of dCas13d-dsRBD-APEX2 localization. Clonal dCas13d-dsRBD-APEX2 (Figure 2C) HEK293T cells expressing untagged hTR along with either hTR gRNA or nontarget (NT) gRNA were fixed and stained with DAPI (nuclei) and anti-V5 antibody to visualize APEX2 (Alexa Fluor 647, green). Scale bar, 10 μm.

**Supplementary Figure 9.**
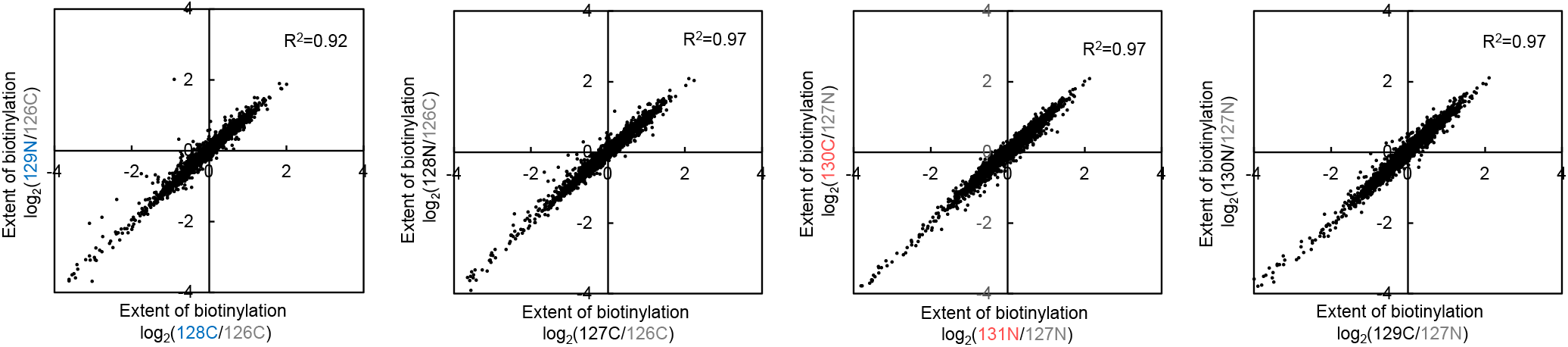
Scatter plots showing the correlation between replicates for both MCP-APEX2 samples and dCas13d-dsRBD-APEX2 samples. The extent of biotinylation by APEX2 for all detected proteins in replicate 1 is plotted against the same values from replicate 2.

**Supplementary Figure 10.**
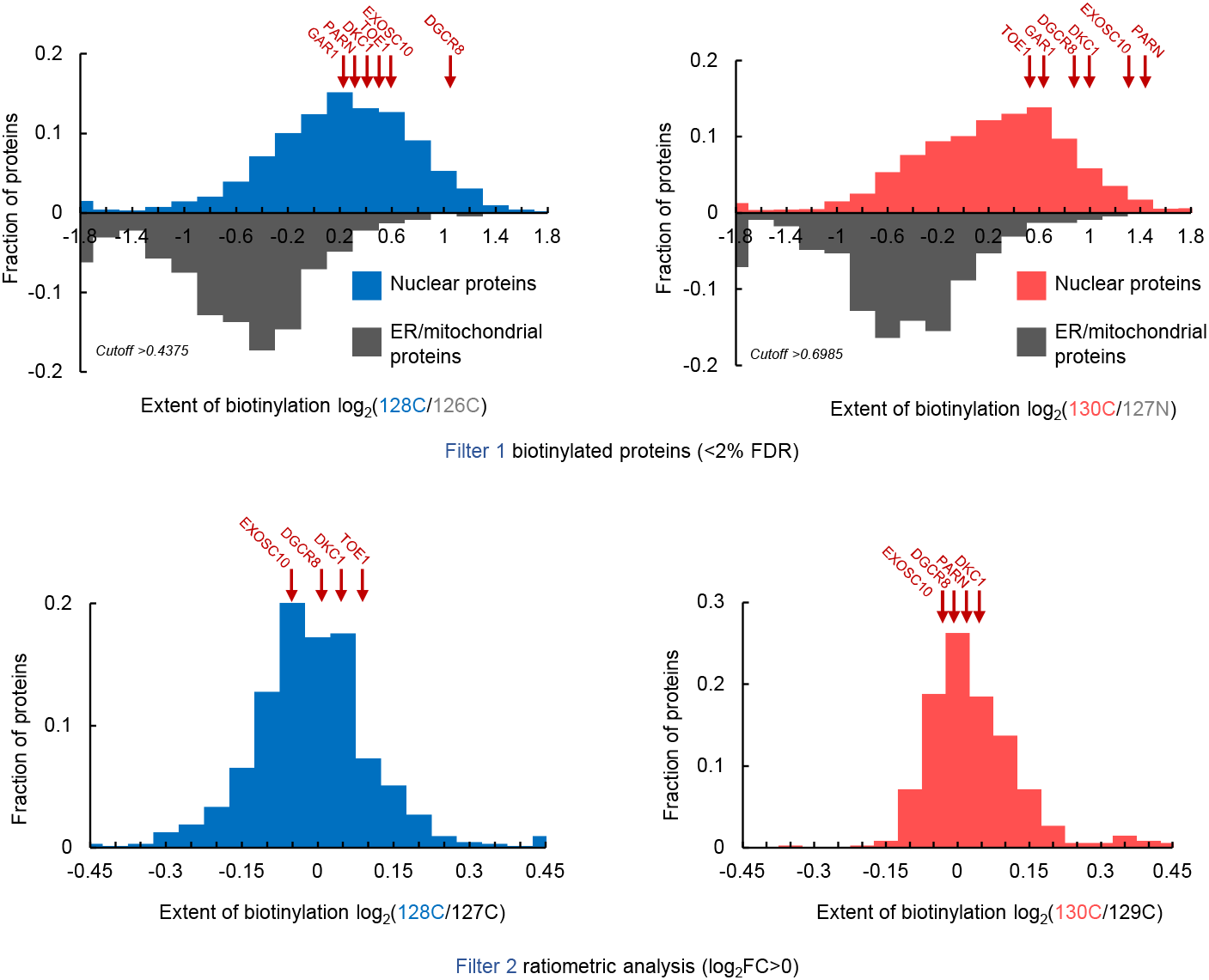
Top: protein density histogram showing how Filter 1 cutoff was determined. True negatives (ER and mitochondrial proteins) are plotted in grey, and nuclear proteins are plotted in blue or red for MCP-APEX2 and dCas13d-dsRBD-APEX2 samples, respectively. Red arrows indicate the TMT ratio of detected hTR binding proteins in Supplementary Table 1. The cutoff was chosen based on <2% false-discovery-rate (FDR) and each replicate is analyzed separately. Bottom: protein density histogram showing how Filter 2 cutoff was determined (after application of Filter 1). Red arrows indicate the TMT ratio of detected hTR binding proteins in Supplementary Table 1. The cutoff was chosen based on preferential enrichment in the targeted APEX2 versus nontargeted APEX2 sample (log_2_ fold change>0). Each replicate was analyzed separately.

**Supplementary Figure 11.**
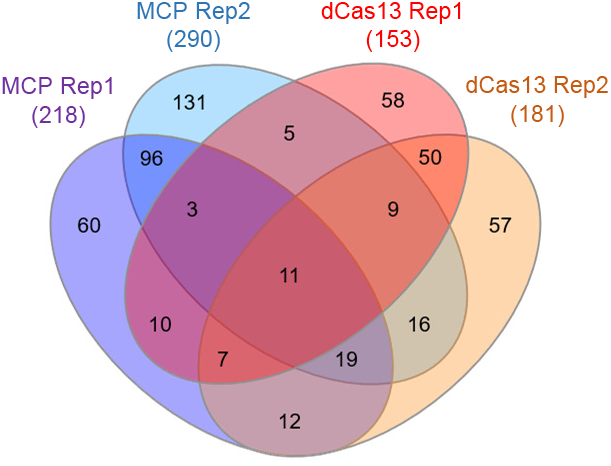
Venn diagram depicting the protein content overlap between proteomic datasets.

**Supplementary Figure 12.**
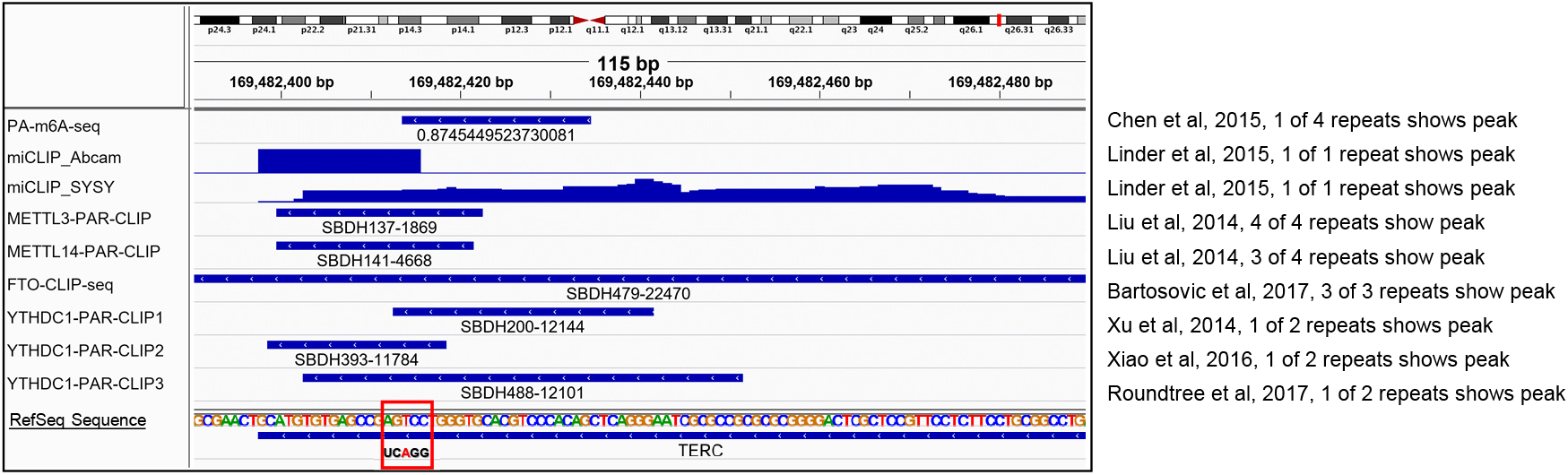
Genome tracks of published sequencing datasets that support m^6^A modification on hTR. The m^6^A consensus motif GGACU is boxed in red with the putative methylated adenosine highlighted.

**Supplementary Figure 13.**
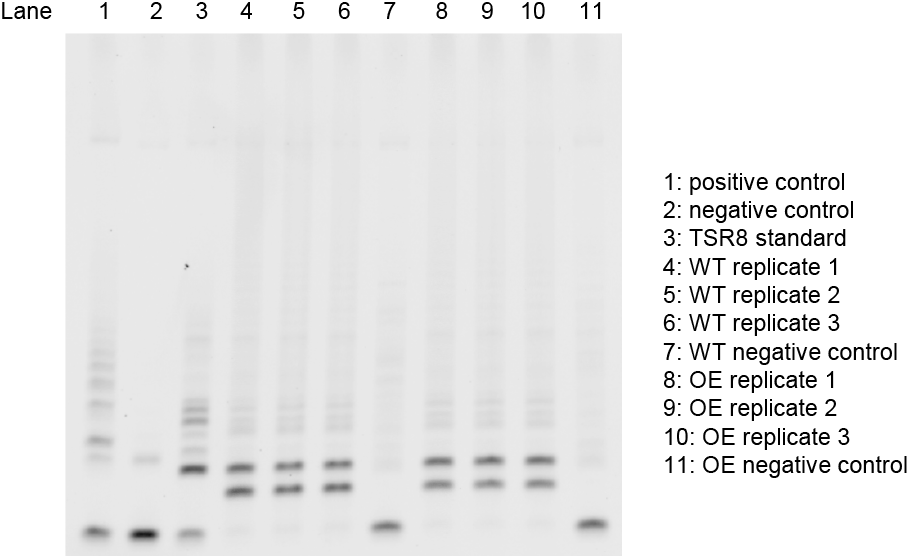
TBE (Tis-Borate-EDTA) gel showing PCR products from the PCR-based telomerase activity assay (TRAPeze kit). Positive control in lane 1 is the extract of telomerase positive cells from the kit; negative control in lane 2 is reaction buffer; TSR8 standard in lane 3 is an oligonucleotide primer that serves as a standard for quantitation; negative controls in lane 7 and 11 are heat-inactivated cell lysates.

## References

1. Müller-Mcnicoll, M. & Neugebauer, K. M. How cells get the message: Dynamic assembly and function of mRNA-protein complexes. Nature Reviews Genetics 14, 275–287 (2013).

2. Lee, S. R. & Lykke-Andersen, J. Emerging roles for ribonucleoprotein modification and remodeling in controlling RNA fate. Trends in Cell Biology 23, 504–510 (2013).

3. Tsai, B. P., Wang, X., Huang, L. & Waterman, M. L. Quantitative profiling of in vivo-assembled RNA-protein complexes using a novel integrated proteomic approach. Mol. Cell. Proteomics 10, (2011).

4. Yoon, J. H., Srikantan, S. & Gorospe, M. MS2-TRAP (MS2-tagged RNA affinity purification): Tagging RNA to identify associated miRNAs. Methods 58, 81–87 (2012).

5. Gong, C., Popp, M. W. L. & Maquat, L. E. Biochemical analysis of long non-coding RNA-containing ribonucleoprotein complexes. Methods 58, 88–93 (2012).

6. Rogell, B. et al. Specific RNP capture with antisense LNA/DNA mixmers. RNA 23, 1290–1302 (2017).

7. Chu, C. et al. Systematic discovery of Xist RNA binding proteins. Cell 161, 404–416 (2015).

8. McHugh, C. A. et al. The Xist lncRNA interacts directly with SHARP to silence transcription through HDAC3. Nature 521, 232–236 (2015).

9. Mili, S., & Steitz, J. A. Evidence for reassociation of RNA-binding proteins after cell lysis: Implications for the interpretation of immunoprecipitation analyses. RNA 10, 1692–1694 (2004).

10. Jazurek, M., Ciesiolka, A., Starega-Roslan, J., Bilinska, K. & Krzyzosiak, W. J. Identifying proteins that bind to specific RNAs - Focus on simple repeat expansion diseases. Nucleic Acids Research 44, 9050–9070 (2016).

11. Faoro, C., & Ataide, S. F. Ribonomic approaches to study the RNA-binding proteome. FEBS Letters 588, 3649–3664 (2014).

12. Ramanathan, M. et al. RNA–protein interaction detection in living cells. Nat. Methods 15, 207–212 (2018).

13. Mukherjee, J. et al. β-Actin mRNA interactome mapping by proximity biotinylation. Proc. Natl. Acad. Sci. U. S. A. 116, 12863–12872 (2019).

14. Han, S., Li, J. & Ting, A. Y. Proximity labeling: spatially resolved proteomic mapping for neurobiology. Current Opinion in Neurobiology 50, 17–23 (2018).

15. Roux, K. J., Kim, D. I., Raida, M. & Burke, B. A promiscuous biotin ligase fusion protein identifies proximal and interacting proteins in mammalian cells. J. Cell Biol. 196, 801–10 (2012).

16. Abudayyeh, O. O. et al. RNA targeting with CRISPR–Cas13. Nature 550, 280–284 (2017).

17. Konermann, S. et al. Transcriptome Engineering with RNA-Targeting Type VI-D CRISPR Effectors. Cell 173, 665–676.e14 (2018).

18. Lam, S. S. et al. Directed evolution of APEX2 for electron microscopy and proximity labeling. Nat. Methods 12, 51–4 (2015).

19. Rhee, H. W. et al. Proteomic Mapping of Mitochondria in Living Cells via Spatially Restricted Enzymatic Tagging. Science (80-.). 339, 1328–1331 (2013).

20. Paek, J. et al. Multidimensional Tracking of GPCR Signaling via Peroxidase-Catalyzed Proximity Labeling. Cell 169, 338–349.e11 (2017).

21. Lobingier, B. T. et al. An Approach to Spatiotemporally Resolve Protein Interaction Networks in Living Cells. Cell 169, 350–360.e12 (2017).

22. Han, S. et al. Proximity Biotinylation as a Method for Mapping Proteins Associated with mtDNA in Living Cells. Cell Chem. Biol. 24, 404–414 (2017).

23. Li, J. et al. Cell-Surface Proteomic Profiling in the Fly Brain Uncovers Wiring Regulators. Cell 180, 373–386.e15 (2020).

24. Theimer, C. A. & Feigon, J. Structure and function of telomerase RNA. Current Opinion in Structural Biology 16, 307–318 (2006).

25. Chen, J. L. & Greider, C. W. Telomerase RNA structure and function: Implications for dyskeratosis congenita. Trends in Biochemical Sciences 29, 183–192 (2004).

26. Zhang, Q., Kim, N. K. & Feigon, J. Architecture of human telomerase RNA. Proc. Natl. Acad. Sci. U. S. A. 108, 20325–20332 (2011).

27. Xi, L., & Cech, T. R. Inventory of telomerase components in human cells reveals multiple subpopulations of hTR and hTERT. Nucleic Acids Res. 42, 8565–8577 (2014).

28. Feng, J. et al. The RNA component of human telomerase. Science (80-.). 269, 1236–1241 (1995).

29. Gazzaniga, F. S. & Blackburn, E. H. An antiapoptotic role for telomerase RNA in human immune cells independent of telomere integrity or telomerase enzymatic activity. Blood 124, 3675–3684 (2014).

30. Kedde, M. et al. Telomerase-independent regulation of ATR by human telomerase RNA. J. Biol. Chem. 281, 40503–40514 (2006).

31. Zheng, G. et al. ALKBH5 Is a Mammalian RNA Demethylase that Impacts RNA Metabolism and Mouse Fertility. Mol. Cell 49, 18–29 (2013).

32. Johansson, H. E. et al. A thermodynamic analysis of the sequence-specific binding of RNA by bacteriophage MS2 coat protein. Proc. Natl. Acad. Sci. U. S. A. 95, 9244–9249 (1998).

33. Abudayyeh, O. O. et al. C2c2 is a single-component programmable RNA-guided RNA-targeting CRISPR effector. Science (80-.). 353, (2016).

34. Fu, D., & Collins, K. Distinct biogenesis pathways for human telomerase RNA and H/ACA small nucleolar RNAs. Mol. Cell 11, 1361–1372 (2003).

35. Jády, B. E., Bertrand, E. & Kiss, T. Human telomerase RNA and box H/ACA scaRNAs share a common Cajal body-specific localization signal. J. Cell Biol. 164, 647–652 (2004).

36. Jády, B. E., Richard, P., Bertrand, E. & Kiss, T. Cell cycle-dependent recruitment of telomerase RNA and cajal bodies to human telomeres. Mol. Biol. Cell 17, 944–954 (2006).

37. Ortega, Á. D., Willers, I. M., Sala, S. & Cuezva, J. M. Human G3BP1 interacts with β-F1-ATPase mRNA and inhibits its translation. J. Cell Sci. 123, 2685–2696 (2010).

38. East-Seletsky, A. et al. Two distinct RNase activities of CRISPR-C2c2 enable guide-RNA processing and RNA detection. Nature 538, 270–273 (2016).

39. Cox, D. B. T. et al. RNA editing with CRISPR-Cas13. Science (80-.). 358, 1019–1027 (2017).

40. Yang, L. Z. et al. Dynamic Imaging of RNA in Living Cells by CRISPR-Cas13 Systems. Mol. Cell 76, 981–997.e7 (2019).

41. Masliah, G., Barraud, P. & Allain, F. H. T. RNA recognition by double-stranded RNA binding domains: A matter of shape and sequence. Cellular and Molecular Life Sciences 70, 1875–1895 (2013).

42. Zhang, C. et al. Structural Basis for the RNA-Guided Ribonuclease Activity of CRISPR-Cas13d. Cell 175, 212–223.e17 (2018).

43. Bevilacqua, P. C. & Cech, T. R. Minor-groove recognition of double-stranded RNA by the double-stranded RNA-binding domain from the RNA-activated protein kinase PKR. Biochemistry 35, 9983–9994 (1996).

44. Eguchi, A. et al. Efficient siRNA delivery into primary cells by a peptide transduction domain-dsRNA binding domain fusion protein. Nat. Biotechnol. 27, 567–571 (2009).

45. Ryter, J. M. & Schultz, S. C. Molecular basis of double-stranded RNA-protein interactions: Structure of a dsRNA-binding domain complexed with dsRNA. EMBO J. 17, 7505–7513 (1998).

46. Chen, J. L., Blasco, M. A. & Greider, C. W. Secondary structure of vertebrate telomerase RNA. Cell 100, 503–14 (2000).

47. Thompson, A. et al. Tandem mass tags: A novel quantification strategy for comparative analysis of complex protein mixtures by MS/MS. Anal. Chem. 75, 1895–1904 (2003).

48. Hung, V. et al. Proteomic mapping of the human mitochondrial intermembrane space in live cells via ratiometric APEX tagging. Mol. Cell 55, 332–41 (2014).

49. Loh, K. H. et al. Proteomic Analysis of Unbounded Cellular Compartments: Synaptic Clefts. Cell 166, 1295–1307.e21 (2016).

50. Mitchell, J. R., Wood, E. & Collins, K. A telomerase component is defective in the human disease dyskeratosis congenita. Nature 402, 551–555 (1999).

51. Macias, S., Cordiner, R. A., Gautier, P., Plass, M. & Cáceres, J. F. DGCR8 Acts as an Adaptor for the Exosome Complex to Degrade Double-Stranded Structured RNAs. Mol. Cell 60, 873–885 (2015).

52. Oughtred, R. et al. The BioGRID interaction database: 2019 update. Nucleic Acids Res. 47, D529–D541 (2019).

53. Khurts, S. et al. Nucleolin interacts with telomerase. J. Biol. Chem. 279, 51508–15 (2004).

54. Hong, J., Lee, J. H. & Chung, I. K. Telomerase activates transcription of cyclin D1 gene through an interaction with NOL1. J. Cell Sci. 129, 1566–79 (2016).

55. Szklarczyk, D. et al. STRING v11: Protein-protein association networks with increased coverage, supporting functional discovery in genome-wide experimental datasets. Nucleic Acids Res. 47, D607–D613 (2019).

56. Gao, X. D. et al. C-BERST: defining subnuclear proteomic landscapes at genomic elements with dCas9–APEX2. Nat. Methods 15, 433–436 (2018).

57. Myers, S. A. et al. Discovery of proteins associated with a predefined genomic locus via dCas9– APEX-mediated proximity labeling. Nat. Methods 15, 437–439 (2018).

58. Benhalevy, D., Anastasakis, D. G. & Hafner, M. Proximity-CLIP provides a snapshot of protein-occupied RNA elements in subcellular compartments. Nat. Methods 15, 1074–1082 (2018).

59. Moon, D. H. et al. Poly(A)-specific ribonuclease (PARN) mediates 3′-end maturation of the telomerase RNA component. Nat. Genet. 47, 1482–1488 (2015).

60. Shukla, S., Schmidt, J. C., Goldfarb, K. C., Cech, T. R. & Parker, R. Inhibition of telomerase RNA decay rescues telomerase deficiency caused by dyskerin or PARN defects. Nat. Struct. Mol. Biol. 23, 286–292 (2016).

61. Deng, T. et al. TOE1 acts as a 3′ exonuclease for telomerase RNA and regulates telomere maintenance. Nucleic Acids Res. 47, 391–405 (2019).

62. Tang, C. et al. ALKBH5-dependent m6A demethylation controls splicing and stability of long 3’-UTR mRNAs in male germ cells. Proc. Natl. Acad. Sci. U. S. A. 115, E325–E333 (2018).

63. Zhang, S. et al. m6A Demethylase ALKBH5 Maintains Tumorigenicity of Glioblastoma Stem-like Cells by Sustaining FOXM1 Expression and Cell Proliferation Program. Cancer Cell 31, 591–606.e6 (2017).

64. Zhang, C. et al. Hypoxia induces the breast cancer stem cell phenotype by HIF-dependent and ALKBH5-mediated m6A-demethylation of NANOG mRNA. Proc. Natl. Acad. Sci. U. S. A. 113, E2047–E2056 (2016).

65. Desrosiers, R., Friderici, K. & Rottman, F. Identification of methylated nucleosides in messenger RNA from Novikoff hepatoma cells. Proc. Natl. Acad. Sci. U. S. A. 71, 3971–3975 (1974).

66. Perry, R. P., Kelley, D. E., Friderici, K. & Rottman, F. The methylated constituents of L cell messenger RNA: Evidence for an unusual cluster at the 5′ terminus. Cell 4, 387–394 (1975).

67. Zhao, B. S., Roundtree, I. A. & He, C. Post-transcriptional gene regulation by mRNA modifications. Nature Reviews Molecular Cell Biology 18, 31–42 (2016).

68. Shi, H., Wei, J. & He, C. Where, When, and How: Context-Dependent Functions of RNA Methylation Writers, Readers, and Erasers. Mol. Cell 74, 640–650 (2019).

69. Baltz, A. G. et al. The mRNA-Bound Proteome and Its Global Occupancy Profile on Protein-Coding Transcripts. Mol. Cell 46, 674–690 (2012).

70. Chen, K. et al. High-resolution N6-methyladenosine (m6A) map using photo-crosslinking-assisted m6A sequencing. Angew. Chemie - Int. Ed. 54, 1587–1590 (2015).

71. Linder, B. et al. Single-nucleotide-resolution mapping of m6A and m6Am throughout the transcriptome. Nat. Methods 12, 767–772 (2015).

72. Liu, J. et al. A METTL3-METTL14 complex mediates mammalian nuclear RNA N6-adenosine methylation. Nat. Chem. Biol. 10, 93–95 (2014).

73. Bartosovic, M. et al. N6-methyladenosine demethylase FTO targets pre-mRNAs and regulates alternative splicing and 3′-end processing. Nucleic Acids Res. 45, 11356–11370 (2017).

74. Xu, C. et al. Structural basis for selective binding of m6A RNA by the YTHDC1 YTH domain. Nat. Chem. Biol. 10, 927–929 (2014).

75. Xiao, W. et al. Nuclear m6A Reader YTHDC1 Regulates mRNA Splicing. Mol. Cell 61, 507–519 (2016).

76. Roundtree, I. A. et al. YTHDC1 mediates nuclear export of N6-methyladenosine methylated mRNAs. Elife 6, (2017).

77. Liu, N. et al. N6-methyladenosine-dependent RNA structural switches regulate RNA-protein interactions. Nature 518, 560–564 (2015).

78. Fazal, F. M. et al. Atlas of Subcellular RNA Localization Revealed by APEX-Seq. Cell 178, 473–490.e26 (2019).

79. Wu, B., Buxbaum, A. R., Katz, Z. B., Yoon, Y. J. & Singer, R. H. Quantifying Protein-mRNA Interactions in Single Live Cells. Cell 162, 211–220 (2015).

80. Wu, B., Eliscovich, C., Yoon, Y. J. & Singer, R. H. Translation dynamics of single mRNAs in live cells and neurons. Science (80-.). 352, 1430–1435 (2016).

81. Grünwald, D. & Singer, R. H. In vivo imaging of labelled endogenous Β-actin mRNA during nucleocytoplasmic transport. Nature 467, 604–607 (2010).

82. Wu, J., Corbett, A. H. & Berland, K. M. The intracellular mobility of nuclear import receptors and NLS cargoes. Biophys. J. 96, 3840–3849 (2009).

83. Kaewsapsak, P., Shechner, D. M., Mallard, W., Rinn, J. L. & Ting, A. Y. Live-cell mapping of organelle-associated RNAs via proximity biotinylation combined with protein-RNA crosslinking. Elife 6, (2017).

84. Li, J. H., Liu, S., Zhou, H., Qu, L. H. & Yang, J. H. StarBase v2.0: Decoding miRNA-ceRNA, miRNA-ncRNA and protein-RNA interaction networks from large-scale CLIP-Seq data. Nucleic Acids Res. 42, D92–7 (2014).

